# Spicing up urinary tract infections: synergistic action of fosfomycin with trans-cinnamaldehyde - insight to the mode of action

**DOI:** 10.64898/2026.06.23.733984

**Authors:** Monika Karczewska, Patryk Strzelecki, Monika Maciąg-Dorszyńska, Małgorzata Kapusta, Agnieszka Pyrczak-Felczykowska, Agnieszka Szalewska-Pałasz, Dariusz Nowicki

**Affiliations:** Department of Bacterial Molecular Genetics, Faculty of Biology, University of Gdansk, Wita Stwosza 59, 80-308, Gdansk, Poland; Bioimaging Laboratory, Faculty of Biology, University of Gdansk, Wita Stwosza 59, 80-308 Gdansk, Poland; Department of Physiology, Medical University of Gdańsk, Gdańsk, Poland

**Author notes:** Corresponding Author Dariusz Nowicki, Department of Bacterial Molecular Genetics, Faculty of Biology, University of Gdansk, Wita Stwosza 59, 80-308, Gdansk, Poland.

**Keywords:** uropathogens, fosfomycin, pyruvate, antimicrobial, G. mellonella

## Abstract

**Objectives:** Fosfomycin (FOS) remains an important therapeutic option for urinary tract infections caused by uropathogenic *Escherichia coli* (UPEC), but specific virulence traits as biofilm formation, metabolic adaptation, and antimicrobial resistance may limit its efficacy. This study investigated whether the natural compound, trans-cinnamaldehyde (t-CA) potentiates FOS activity against UPEC and explored the underlying mechanisms of its effect

**Methods:** The interaction between t-CA and FOS was assessed using checkerboard assays, time-kill analysis. We evaluated biofilm viability and structure using confocal and scanning microscopy as well as catheter-associated biofilm models. Next, effects on membrane integrity, cell-surface properties, membrane potential, intracellular pyruvate levels, and resistance evolution during serial passage were evaluated. Molecular docking was used to explore potential interactions of t-CA with enzymes involved in pyruvate metabolism. *Galleria mellonella* infection model was employed to evaluate *in vivo* therapeutical efficiency.

**Results:** t-CA potentiated FOS activity against laboratory, reference, and clinical UPEC strains, with synergistic or additive interactions observed across the tested collection. The combination enhanced bacterial killing, reduced biofilm viability and biomass, and disrupted biofilm architecture. In catheter-associated biofilms, combined treatment markedly impaired surface-associated UPEC communities. t-CA reduced extracellular matrix abundance and altered cell-surface hydrophobicity and membrane potential without inducing detectable oxidative stress. Mechanistically, t-CA affected pyruvate homeostasis, reduced intracellular pyruvate levels, and phenotypically intersected with the BtsSR pyruvate-sensing pathway. Serial exposure to FOS alone rapidly increased MIC, whereas t-CA limited this phenomenon and did not itself promote reduced susceptibility. The compounds combination also improved survival of UTI89-infected *G. mellonella* larvae.

**Conclusions:** t-CA enhances FOS activity against UPEC through complementing the antibiofilm and metabolic effects. By weakening biofilm matrix integrity, perturbing pyruvate homeostasis, and limiting FOS-associated MIC elevation, t-CA represents a promising adjuvant candidate for improving FOS efficacy against biofilm-associated UPEC infections.

## 1. Introduction

Antimicrobial resistance (AMR) is accelerating worldwide and increasingly undermines the clinical utility of first-line antibiotics, creating an urgent need for strategies that restore or potentiate antibacterial activity while limiting further resistance selection. Recent surveillance data indicate that resistance is already widespread among pathogens responsible for common community- and hospital-acquired infections, with particularly high rates reported for urinary tract and bloodstream isolates in several regions(WHO, 2025). The long-term outlook remains alarming. Influential global assessments have projected that, without major intervention, AMR could rise to a scale comparable to (or exceeding) cancer mortality by mid-century, reflecting both increasing incidence of hard-to-treat infections and limited replenishment of the antibiotic pipeline.

Urinary tract infections (UTIs) are among the most prevalent bacterial infections and a major driver of antibiotic consumption in outpatient care (WHO, 2025). Uropathogenic *Escherichia coli* (UPEC) dominates uncomplicated UTIs, but treatment is increasingly complicated by resistance to commonly used oral agents. Surveillance programs and regional studies consistently report high resistance rates among outpatient uropathogens to commonly prescribed antibiotics, including trimethoprim–sulfamethoxazole, fluoroquinolones, and β-lactams. In some settings, resistance exceeds 30% for selected drug classes, emphasizing the need for therapeutic strategies that retain activity against resistant and multidrug-resistant isolates. Although UPEC infections are generally associated with low mortality, in otherwise healthy individuals, they have significantly more serious impact on elderly or immunocompromised patients leading to e.g. life threatening urosepsis (Mancuso *et al*., 2023). For the public health, notably, these strains constitute an important reservoir and vehicle for the dissemination of antimicrobial resistance determinants (Karczewska *et al*., 2023; Rahman *et al*., 2022). Moreover, the recurrent nature of UTIs often requires repeated or prolonged antibiotic therapy, resulting in a substantial cumulative economic burden for both patients and healthcare systems (Iskandar *et al*., 2021; Karczewska *et al*., 2023).

Fosfomycin (FOS) remains a valuable option in the UTIs treatment strategy (Figure 1A)(Falagas *et al*., 2010). Clinically, a single 3-g oral dose of fosfomycin trometamol is recommended as a first-line treatment for uncomplicated lower UTIs in women in multiple guidelines and regulatory evaluations (Zheng *et al*., 2022). Mechanistically, FOS is a phosphoenolpyruvate (PEP) analogue that inhibits the first committed step of peptidoglycan biosynthesis by covalently targeting MurA (UDP-GlcNAc enolpyruvyl transferase), an essential enzyme for cell wall formation. In addition to its broad activity against many uropathogens, including ESBL-producing *E. coli*, FOS benefits from favorable urinary exposure and a comparatively low propensity for widespread resistance in many regions-although resistance can emerge and is often linked to altered uptake (e.g., transporter function) or enzymatic inactivation (Mattioni Marchetti *et al*., 2023).

**Figure 1.**
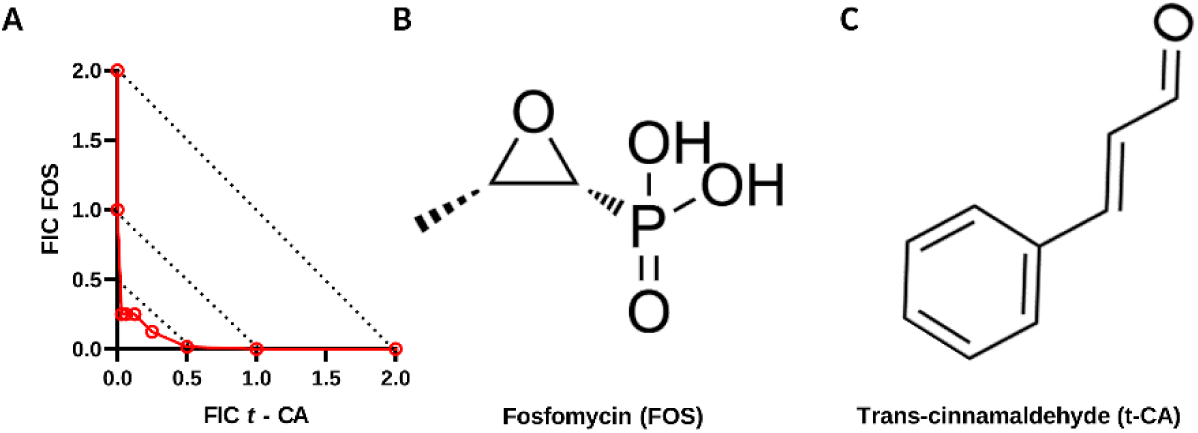
Fosfomycin/trans-cinnamaldehyde interaction and chemical structures of the tested compounds. (A) Representative isobologram of the interaction between fosfomycin (FOS) and trans-cinnamaldehyde (t-CA) against *E. coli* MG1655. Fractional inhibitory concentration (FIC) values are shown for FOS and t-CA, and dotted diagonal lines indicate the thresholds used for fractional inhibitory concentration index (FICI) classification. The red interaction curve falls within the synergistic range, indicating potentiation of FOS activity by t-CA. (B) Chemical structure of FOS, a MurA-targeting antibiotic that inhibits peptidoglycan biosynthesis. (C) Chemical structure of t-CA, a plant-derived phenylpropanoid evaluated as a phytochemical adjuvant.

A key concept that has gained traction in recent years is that FOS efficacy is tightly coupled to bacterial physiology, particularly metabolic state and transport capacity (Gil-Gil & Martinez, 2022, 2024). Because FOS relies on active transport into the cytoplasm (notably via systems influenced by central carbon metabolism), metabolic perturbations can alter intracellular drug accumulation and thereby modulate susceptibility. Consistent with this, recent systems-level work demonstrates that metabolic state can shape FOS activity and may even promote the emergence of FOS resistance under certain perturbations, reinforcing the idea that metabolic context is a decisive layer of antibiotic action beyond the canonical target-inhibitor interaction (Peng *et al*., 2025; Verhulsdonk *et al*., 2026). This physiology dependence suggests a practical opportunity: adjunct compounds that reprogram metabolism or stress responses may sensitize bacteria to FOS, lowering the effective concentration required for growth inhibition and potentially slowing resistance selection during repeated exposure.

Natural products and plant-derived small molecules represent a valuable source of antibiotic adjuvants because their biological activity often extends beyond direct growth inhibition and includes modulation of stress adaptation, virulence, and central metabolic pathways (Groleau *et al*., 2026; Strzelecki *et al*., 2025). Among these compounds, trans-cinnamaldehyde (t-CA, Figure 1B) is particularly relevant due to its ability to affect several interconnected aspects of bacterial physiology. Our previous work demonstrated that t-CA is active against pathogenic *E. coli* and that its antibacterial effect is accompanied by attenuation of virulence-associated processes rather than being explained solely by nonspecific membrane damage(Karczewska *et al*., 2024). More recent transcriptomic and biochemical analyses showed that sub-inhibitory t-CA concentration induces RelA-dependent accumulation of (p)ppGpp and extensive transcriptional reprogramming, including repression of iron-acquisition systems, quorum-sensing-associated pathways, and acid-resistance mechanisms (Karczewska *et al*., 2026). These changes were substantially weaker in the absence of RelA, indicating that activation of the stringent response is a central component of the cellular response to t-CA. In parallel, t-CA impaired extracellular matrix production and reduced biofilm biomass and metabolic activity, confirming that its effects extend to physiologically heterogeneous bacterial populations and affects global metabolic pathways. Thus, t-CA should not be considered merely as a membrane-active phytochemical, but rather as a stress- and metabolism-modulating compound capable of altering the physiological state that determines antibiotic susceptibility. This multi-target activity provides a mechanistic basis for testing t-CA as an adjuvant to FOS, whose uptake and bactericidal activity are strongly influenced by bacterial metabolic status and active cell-wall synthesis. We therefore reasoned that t-CA-mediated disruption of stress adaptation and metabolic homeostasis could increase the susceptibility of *E. coli* to FOS and reduce the concentrations required for effective growth inhibition.

In this study, we investigated whether combining FOS with t-CA enhances inhibitory activity against *E. coli* and whether such enhancement is consistent with a metabolism-linked interaction. We hypothesized that these agents can act synergistically through interconnected physiological pathways whereby t-CA-driven metabolic and/or stress perturbations sensitize *E. coli* to MurA inhibition by FOS leading to stronger growth suppression at reduced concentrations compared with monotherapy. In addition, given the known propensity of bacterial populations to adapt under sustained antibiotic pressure, we examined whether the combination influences the trajectory of susceptibility during serial exposure, a feature that would further support the value of metabolic adjuvants as resistance-mitigating partners for existing first-line drugs.

## 2. Materials and methods

### 2.1 Bacterial strains and growing conditions

Bacterial strains used in this study are listed in Table X. Strains were maintained as glycerol stocks at −80 °C and, prior to each experiment, were streaked onto solid agar (LB Miller, Sigma) plates and incubated under the appropriate conditions, typically at 37 °C. Where indicated, strains were grown on Mueller–Hinton II (MH II, Merck) or YESCA broth (1 g/L yeast extract and 10 g/L casamino acids). Single colonies were selected to initiate overnight cultures in the relevant broth medium and then used for the appropriate assays. Where indicated, cells were washed with sterile buffered saline (PBS).

### 2.2 Susceptibility testing

Susceptibility testing was conducted according to standard CLSI methods (CLSI, 2018). Briefly, the microdilution assay was employed to establish minimum inhibitory concentrations (MICs) of the tested compounds against indicated bacterial strains. The interaction between the antibiotics and *t*-CA was subsequently evaluated using the checkerboard method to determine the fractional inhibitory concentration index (FICI). The assay was performed in 96-well microtiter plates using the previously determined MIC values as a baseline. Stock solutions of the compounds were prepared at 16 times the respective MIC, followed by serial two-fold dilutions of each drug. The first compound was diluted along the ordinate (rows), while the second was diluted along the abscissa (columns), after which the solutions were combined. Each microtiter well was then inoculated with 100 µL of bacterial inoculum to achieve a final concentration of 5 × 10^^5^ CFU/mL at 200 µL of broth. The plates were incubated at 37°C for 20 hours, and the absorbance at 570 nm was measured using an EnSpire plate reader (PerkinElmer, Singapore). FICI was calculated as:

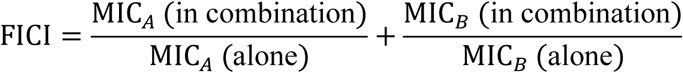

Interactions were interpreted as synergistic (FICI ≤ 0.5), additive (0.5 < FICI ≤ 1), indifferent (1 < FICI ≤ 4), or antagonistic (FICI > 4)

Bacterial kill kinetics were assessed using a time-kill assay (Klepser *et al*., 1998). Briefly, an fresh bacterial culture was adjusted to 0.5 McFarland and diluted in PBS to an initial inoculum of approximately 1 × 10^5 CFU/mL. The suspension was exposed to the indicated antimicrobial agents at the specified concentrations and incubated at 37 °C. At indicated time points, aliquots were withdrawn, serially diluted in sterile PBS, and plated on agar for CFU enumeration after overnight incubation at 37 °C. Bactericidal activity was expressed as log^10^(CFU/mL) over time and presented as time-kill curves.

### 2.3 Live/dead flow cytometry analysis

Live/dead analysis was performed using a two-color fluorescence assay (Thermo Fisher Scientific; L7012) (Konieczny *et al*., 2021). Overnight cultures of *E. coli* strains were prepared and adjusted to approximately 0.5 McFarland. Cells were then exposed to t-CA and fosfomycin at the indicated concentrations and incubated at 37 °C by 120min. Following treatment, cells were stained with a SYTO dye and propidium iodide (PI). SYTO labels all cells, whereas PI penetrates only cells with compromised membranes, enabling discrimination of live (SYTO-positive/PI-negative) and membrane-compromised/dead (PI-positive) populations. Samples were acquired on an Amnis FlowSight imaging flow cytometer (Luminex). Single cells were gated based on brightfield features to exclude aggregates and debris. Fluorescence compensation was performed using single-stained controls. SYTO fluorescence was excited with the 488 nm laser and collected in the green channel, whereas PI fluorescence was collected in the red channel. Data were analyzed using IDEAS (Luminex) and FlowJo. Live and dead populations were quantified using fluorescence thresholds defined from appropriate controls.

### 2.4 Antimicrobial activity evaluation in biofilms

The catheter biofilm assay was performed based on Borowicz et al. with minor modifications (Borowicz *et al*., 2023). Bacteria were grown overnight at 25 °C in YESCA medium (1 g/L yeast extract and 10 g/L casamino acids) with shaking (120 rpm), adjusted to 4 McFarland units, and diluted 1:100 in fresh YESCA. Sterile Nelaton catheters (Unomedical, UK) were cut into 10 cm segments and connected to sterile syringes. The bacterial suspension was drawn into each catheter-syringe assembly (final volume ≤ 3 mL) and sealed aseptically. Catheters were incubated statically at 25 °C for 72 h to allow biofilm development. To compare antibiofilm activity, catheters were incubated with: (i) inoculum without additives (positive biofilm control), (ii) inoculum supplemented with FOS at the indicated concentration, (iii) inoculum supplemented with t-CA at the indicated concentration, and a combined-treatment condition (FOS+t-CA; MIX) was included. A sterility control consisted of catheters filled with sterile medium only. After incubation, catheters were opened and gently rinsed with 20 mL distilled water to remove planktonic cells. Biofilm biomass was quantified by staining with 1% (w/v) crystal violet for 20 min at room temperature, rinsing until the wash was clear, photographing the catheters, and solubilizing the bound dye with 2 mL of 33% (v/v) acetic acid; absorbance was measured at 570 nm (BioSpectrometer, Eppendorf, Germany). In parallel, biofilm viability was assessed using an MTT reduction assay: after rinsing, catheter segments were incubated with MTT in fresh medium in the dark, the resulting formazan was dissolved in an organic solvent, and absorbance was measured spectrophotometrically (typically at 570 nm) and expressed relative to the untreated biofilm control after background subtraction (sterility control).

### 2.5 Microscopy techniques evaluation

Fluorescence staining was routinely assessed by fluorescence microscopy using Leica DMI4000B inverted microscope fitted with a DFC365FX camera (Leica Microsystems, Germany). Scanning electron microscopy (SEM) was employed for imagining of biofilm structure morphology in medical devices. Catheter segments with biofilm were aseptically cut and gently rinsed with sterile PBS to remove loosely attached cells and debris. Samples were fixed in 2.5% (v/v) glutaraldehyde in PBS for 1–2 h at room temperature under gentle agitation, then rinsed three times with PBS. Post-fixation was performed in 1% (w/v) osmium tetroxide (OsO₄) for 1–2 h at room temperature, followed by three rinses with sterile distilled water. Specimens were dehydrated through a graded ethanol series (30%, 50%, 70%, 90%, and 100% ethanol; 10–15 min each step), using fresh ethanol for each step and ensuring complete dehydration. Samples were then dried by critical point drying (CPD), mounted onto aluminum SEM stubs, and sputter-coated with gold using a Spi Module Sputter Coater. Imaging was performed using a Philips XL-30 (FEI) scanning electron microscope (Laboratory of Bioimaging, University of Gdańsk, Gdańsk, Poland) operated at an accelerating voltage of 5 or 15 kV.

Confocal laser scanning microscopy (CLSM). Biofilms of UTI89 were grown in LabTek™ chamber slides as described in Sect. 2.8. After treatment (FOS 2×MIC, t-CA 2×MIC, or combination: FOS 1×MIC + t-CA 1×MIC) and washing, biofilms were stained using the LIVE/DEAD BacLight Bacterial Viability Kit (Thermo Fisher Scientific; L7012). The staining solution was prepared by adding 1.5 µL SYTO 9 and 1.5 µL propidium iodide (PI) to 1 mL saline, followed by incubation at room temperature for 20 min protected from light. The staining mixture was removed, chambers were detached from the slide, and samples were covered with a coverslip and mounted using low-melting agarose. Bacterial biofilms were imaged using a Leica Stellaris 5 confocal laser scanning microscope controlled with LAS X software. Image stacks were acquired in xyz mode using an HC PL FLUOTAR L 40×/0.60 dry objective, with a zoom factor of 1. The pinhole was set to 1 Airy unit. Line averaging was set to 3, while frame averaging, frame accumulation and line accumulation were set to 1. Excitation was provided by a white light laser (WLL), and two acquisition settings were used. In the first setting, excitation was performed at 495 nm with a laser line intensity of 2.29%, and detector gain and offset were set to 25.8 and 0, respectively. In the second setting, excitation was performed at 549 nm with a laser line intensity of 5.41%, and detector gain and offset were set to 63.5 and 0, respectively. For each analysed biofilm, confocal z-stacks consisting of nine optical sections were acquired. Orthogonal views were generated from the original z-stacks. The z-stack datasets were subsequently transferred to LAS AF software for three-dimensional visualisation and reconstruction. Comparable samples were imaged using the same acquisition settings.

### 2.6 Resistance evolution assay

Evolution experiments were performed by serial passaging under antibiotic selection. Overnight cultures were initiated from fresh grown single colonies and resuspended to OD_600_ = 0.005. Cultures were used to inoculate 96-well plates containing LB and a two-fold serial dilution gradient of fosfomycin (with or without t-CA at the indicated fixed concentration, where applicable) and incubated for 24 h at 37 °C with agitation. Optical density was measured using an EnSpire microplate reader. Wells were considered to show growth if OD reached at least 50% of the no-drug LB control. For each passage, the culture growing at the highest fosfomycin concentration was transferred into fresh LB containing the corresponding two-fold fosfomycin gradient (± t-CA) and adjusted again to OD600 = 0.005. Five serial passages were performed unless resistance reached the highest fosfomycin concentration soluble in LB. In control experiments performed with t-CA alone (at the same fixed concentration as in combination assays), no increase in tolerance was observed under the applied conditions. Experiment was done at least three times in triplicates

### 2.8 Molecular docking

Molecular docking of t-CA was performed using CB-Dock2 with automatic pocket detection and docking (Liu *et al*., 2022; Yang *et al*., 2022). The t-CA ligand (CAS: 14371-10-9) was provided as a 3D structure file (MolFile). Receptor structures were obtained from the Protein Data Bank: AceE (PDH E1; PDB ID: 2IEA), AceF (PDH E2; PDB ID: 8OSY), LpdA (PDH E3; PDB ID: 4JQ9), and PoxB (PDB ID: 3EYA). Prior to docking, co-crystallized ligands and all crystallographic water molecules were removed from each receptor structure. Docking poses were ranked according to the CB-Dock2 scoring output. Protein-ligand interactions and 2D contact residues were analyzed and illustrated using BIOVIA Discovery Studio Visualizer. Docking poses were prioritized based on their localization within the predicted binding pockets corresponding to catalytically relevant regions (active-site) of each target, as inferred from pocket placement relative to functionally important residues and the overall interaction pattern (hydrogen bonding, π-interactions, and hydrophobic contacts).

### 2.9 Cell-surface alteration evaluation

Cell surface hydrophobicity was assessed by using the MATH assay (Pembrey *et al*., 1999). Briefly, overnight bacterial cultures were refreshed and grown to an optical density of OD = 0.2. Cultures were then exposed to t-CA, FOS, or their combination (MIX), while untreated cells served as the control. Samples were incubated with shaking, and aliquots were collected after 30 and 60 min of treatment. The absorbance of each bacterial suspension was measured at 550 nm and recorded as A₀. Xylene was then added to each sample, and the mixtures were vigorously vortexed to allow interaction between bacterial cells and the hydrocarbon phase. Samples were left undisturbed until complete phase separation. The lower aqueous phase was carefully collected, and its absorbance was measured again at 550 nm and recorded as A₁. Cell-surface hydrophobicity was calculated as the percentage of cells adhering to xylene according to the formula:

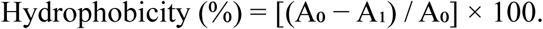

Membrane potential effects were conducted using the voltage-sensitive dye 3,3’-dipropylthiadicarbocyanine iodide [DiSC_3_(5)] (Buttress *et al*., 2022). Bacterial cultures were grown to an optical density of OD = 0.5 and centrifuged for 15 min at 4000 rpm. The cell pellet was resuspended in PBS containing 1% DMSO. The cell suspension was transferred to wells of a black microplate and mixed with DiSC₃(5), prepared in DMSO, to a final concentration of 1 µM. Samples were incubated for 25 min at 37°C in the dark to allow dye equilibration. Fluorescence was then monitored at an excitation wavelength of 652 nm and an emission wavelength of 671 nm at 1-min intervals until a stable baseline signal was obtained. After baseline stabilization, the tested compounds were added, and fluorescence measurement was continued at 1-min intervals. Increased fluorescence was interpreted as a change in membrane potential consistent with membrane depolarization.

### 2.10 Antimicrobial efficiency of the antimicrobial agents in *G. mellonella in vivo* model

Greater wax moth larvae *Galleria mellonella* L. (Lepidoptera: Pyralidae) (Exoticroom, Łódź, Poland) were stored in the dark at 20 °C until use. Healthy larvae weighing approximately 250–300 mg, without signs of melanization or visible dark spots on the cuticle, were selected for experiments (Budziaszek *et al*., 2023). Larvae were infected with *E. coli* UTI89 at a dose of 10^5^ CFU per larva using a Hamilton syringe fitted with a blunt-ended needle. The bacterial suspension (10 µL) was injected into the last left proleg. One hour post-infection, larvae received a second injection (10 µL) into the last right proleg containing either t-CA and FOS in combination or each compound alone. Control larvae were injected with phosphate-buffered saline (PBS). Following treatment, larvae were placed in Petri dishes lined with Whatman filter paper and incubated at 37 °C in the dark. Survival and health status were monitored every 24 h for 4 consecutive days, with morbidity assessed by changes in behavior and melanization. Larvae were considered dead upon lack of movement in response to touch and the presence of extensive dark pigmentation. Each treatment group contained 15 larvae, and all experiments were performed at least three independent times.

### 2.11 Statistic methods

All experiments were performed using at least three independent biological replicates, unless stated otherwise. Data are presented as the mean ± standard deviation (SD), as specified in the corresponding figure legends. Comparisons between two groups were performed using an unpaired, two-tailed Student’s t-test. Comparisons involving three or more groups were analyzed by one-way analysis of variance (ANOVA), followed by an appropriate multiple-comparisons post hoc test, as indicated in the figure legends. Statistical analyses were performed using GraphPad Prism version 9.0 (GraphPad Software, San Diego, CA, USA). Differences were considered statistically significant at *p* < 0.05.

## 3. Results

### Susceptibility and interactions of antimicrobial compounds

To identify antibiotic combinations capable of enhancing the antibacterial activity of t-CA, we first screened its interactions with representatives of several antibiotic classes using the checkerboard assay (Figure S1). Among the tested compounds, FOS produced the most pronounced and reproducible synergistic interaction with t-CA (Figure 1C), whereas the remaining antibiotics showed predominantly additive, or even adverse effect, observed for oxytetracycline.

To determine whether t-CA enhances the antibacterial activity of FOS, their interaction was evaluated against the laboratory strain MG1655, the reference UPEC strains UTI89 and CFT073, and seven clinical *E. coli* isolates. The MIC of t-CA alone was 500 mg/L for all tested strains, whereas the MIC of FOS alone ranged from 2 to 4 mg/L. In combination, the effective concentrations of both compounds were substantially reduced.

Synergistic interactions were observed for eight of the ten tested strains, with FICI values ranging from 0.28125 to 0.500. The strongest interaction was detected for UTI89, for which the effective concentration of t-CA decreased from 500 to 16 mg/L and that of FOS from 4 to 1 mg/L, resulting in a FICI of 0.28125. This corresponded to approximately a 31-fold reduction in the effective t-CA concentration and a four-fold reduction in the effective FOS concentration compared with the respective monotherapies. A similarly strong interaction was observed for KB082, with a FICI of 0.3125.

Checkerboard survival maps further demonstrated that the combinations associated with FICI values below 0.5 coincided with marked reductions in bacterial survival (Figure 2A). The strongest reduction was observed when both compounds were used at fractional concentrations below their respective MICs, indicating that the interaction was not explained by the independent activity of either compound alone.

**Figure 2.**
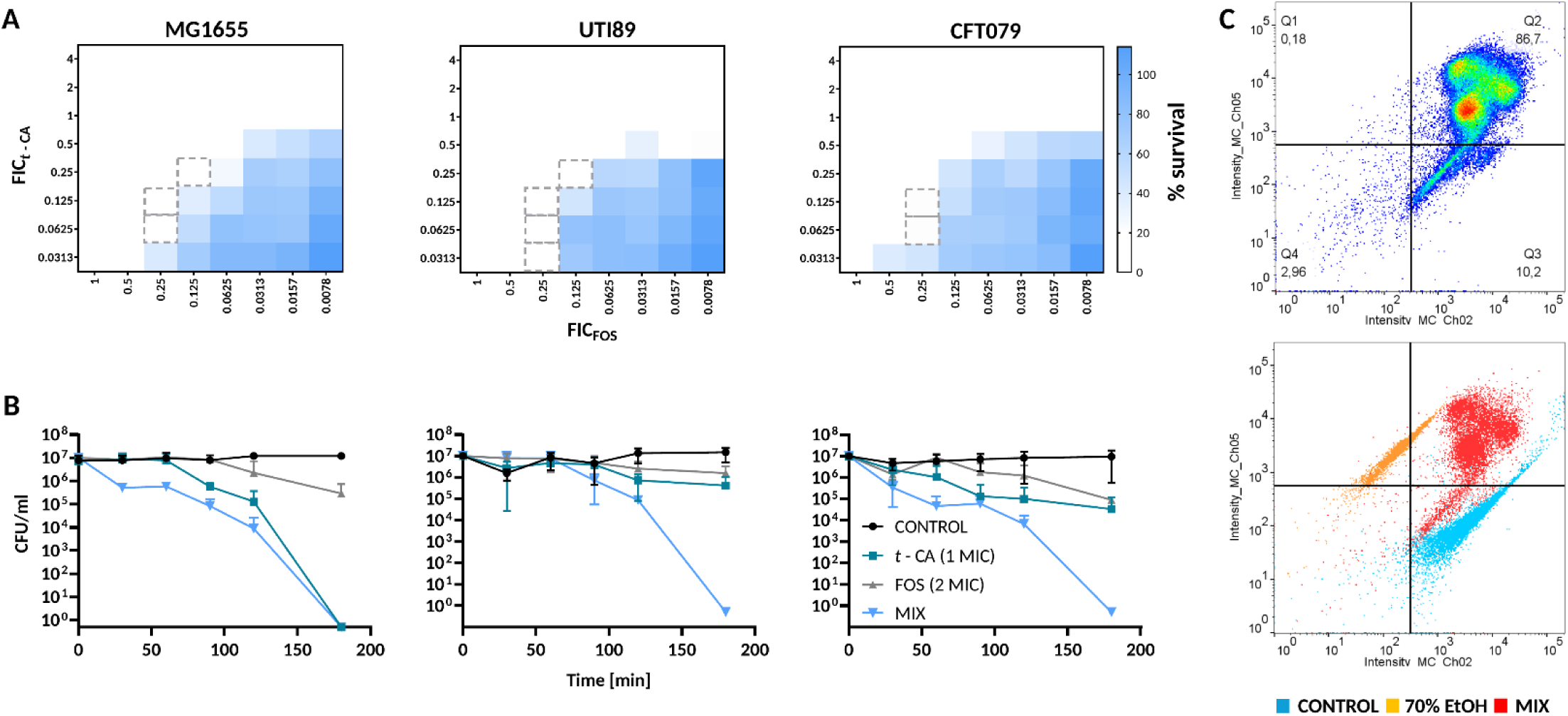
Trans-cinnamaldehyde potentiates the antibacterial activity of fosfomycin against *Escherichia coli*. (A) Checkerboard survival maps showing the interaction between fosfomycin (FOS) and trans-cinnamaldehyde (t-CA) against *E. coli* MG1655 and the uropathogenic *E. coli* (UPEC) strains UTI89 and CFT073. The x-axis represents the fractional inhibitory concentration (FIC) of FOS, whereas the y-axis represents the FIC of t-CA. Color intensity indicates bacterial survival, and dashed rectangles mark concentration combinations corresponding to synergistic interactions. (B) Time-kill kinetics of MG1655, UTI89, and CFT073 exposed to t-CA, FOS, or their combination. Bacterial survival was monitored over 180 min and expressed as colony-forming units per milliliter (CFU/mL). (C) Flow cytometric analysis of membrane integrity after treatment (120 min) with the FOS/t-CA combination (0.5 x 0.5 MIC). UTI89 cells were stained with SYTO 9 and propidium iodide (PI). Untreated live cells and ethanol-killed cells were used as controls. The combination (MIX) shifted the bacterial population toward the PI-positive fraction, indicating extensive membrane damage and loss of viability.

The synergistic effect was further validated in time-kill experiments (Figure 2B). In all three strains, the combination produced a faster and more extensive decline in viable cell counts than either monotherapy. In UTI89 and CFT073, treatment with FOS or t-CA alone resulted only in partial reductions in CFU/mL, whereas the combination decreased bacterial counts to the detection limit after 180 min. A similar effect was observed for MG1655, although t-CA alone displayed stronger intrinsic activity against this strain than against the UPEC isolates. Nevertheless, the combination was consistently more bactericidal than FOS alone and was required for complete killing of the tested UPEC and wild-type strains under the applied conditions.

The effect of the combination on membrane integrity was examined using SYTO/PI staining followed by imaging flow cytometry (Figure S2, 2C). Treatment with the combination caused a marked shift from the live-cell population toward the PI-positive fraction. Approximately 86.7% of cells were classified as membrane-compromised or dead, whereas only 10.2% retained the fluorescence profile characteristic of viable cells. The distribution of the treated population partially overlapped with that of the 70% ethanol control and was clearly separated from the untreated control. These results indicate that the enhanced bactericidal activity of the t-CA/FOS combination is accompanied by extensive loss of membrane integrity.

### Combining the t-CA and FOS agents reduces biofilm viability and disrupts its architecture

The effect of t-CA/FOS combination on biofilm formation was first screened in a 96-well plate model (Figure 3). Fluorescence imaging after SYTO 9/PI staining showed that untreated biofilms were dominated by SYTO 9-positive cells, whereas exposure to either t-CA or FOS increased the proportion of PI-positive cells. The most pronounced shift toward the PI-positive population was observed following treatment with the combination, indicating extensive membrane damage and loss of viability within the biofilm (Figure 3A). This observation was confirmed by the MTT assay. Both compounds applied individually reduced the metabolic activity of biofilm-associated cells compared with the untreated control, although the magnitude of the effect differed between strains (Figure 3B). In all three tested strains, the combination produced the strongest decrease in MTT reduction. The reduction was particularly marked for MG1655 and CFT073, while a similar tendency was observed for UTI89. These results indicate that the combined treatment impairs biofilm-associated cell viability more effectively than either compound used alone.

**Figure 3.**
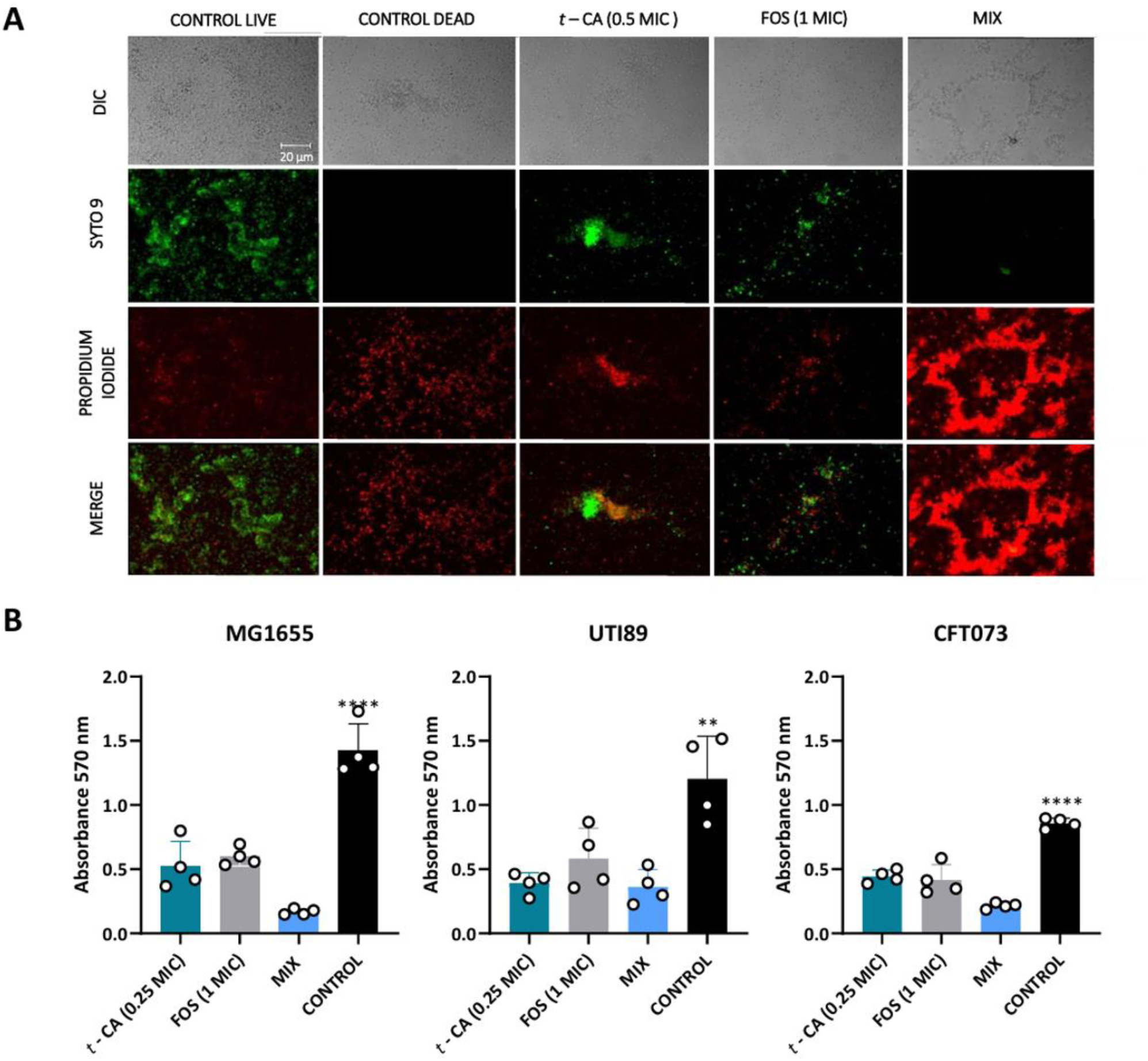
Trans-cinnamaldehyde enhances the antibiofilm activity of fosfomycin in a 96-well plate biofilm model. (A) Representative fluorescence microscopy images of UTI89 biofilms stained with SYTO 9 and propidium iodide (PI) after treatment with trans-cinnamaldehyde (t-CA; 0.5×MIC), fosfomycin (FOS; 1×MIC), or their combination (MIX). Untreated live biofilms and ethanol-killed biofilms were included as staining controls. DIC, SYTO 9, PI, and merged channels are shown. SYTO 9-positive cells are shown in green, whereas PI-positive cells are shown in red and indicate membrane-compromised or dead cells. Scale bar: 20 µm. (B) Metabolic activity of biofilms formed by *Escherichia coli* MG1655 and the uropathogenic *E. coli* strains UTI89 and CFT073, determined using the MTT reduction assay. Biofilms were exposed to t-CA, FOS, or MIX, and absorbance was measured at 570 nm. Data are presented as mean ± SD from independent biological replicates. Statistical significance was assessed relative to the untreated control; **p < 0.01, ***p < 0.001, ****p < 0.0001.

The structural effects of the treatment were subsequently examined by confocal microscopy and in a catheter-associated biofilm model (Figure 4). Confocal imaging showed that FOS and t-CA reduced biofilm continuity and increased the proportion of PI-positive cells, while the combination caused the most pronounced disruption of three-dimensional organization, yielding a thinner and highly fragmented biofilm dominated by membrane-compromised cells (Figure 4A). These changes were evident relative to the untreated control, which formed a dense and predominantly viable biofilm. Even more pronounced pattern was observed on catheter surfaces (Figure 4B). FOS treatment reduced bacterial accumulation and the overall extent of surface colonization, whereas t-CA caused a pronounced depletion of the matrix surrounding the cells and disrupted the cohesion of bacterial aggregates. The combination further enhanced both effects, producing the lowest bacterial coverage and the most substantial loss of matrix-associated material. Quantitative measurements confirmed that the combined treatment caused the greatest reduction in both biofilm viability and total biomass (Figure 4C).

**Figure 4.**
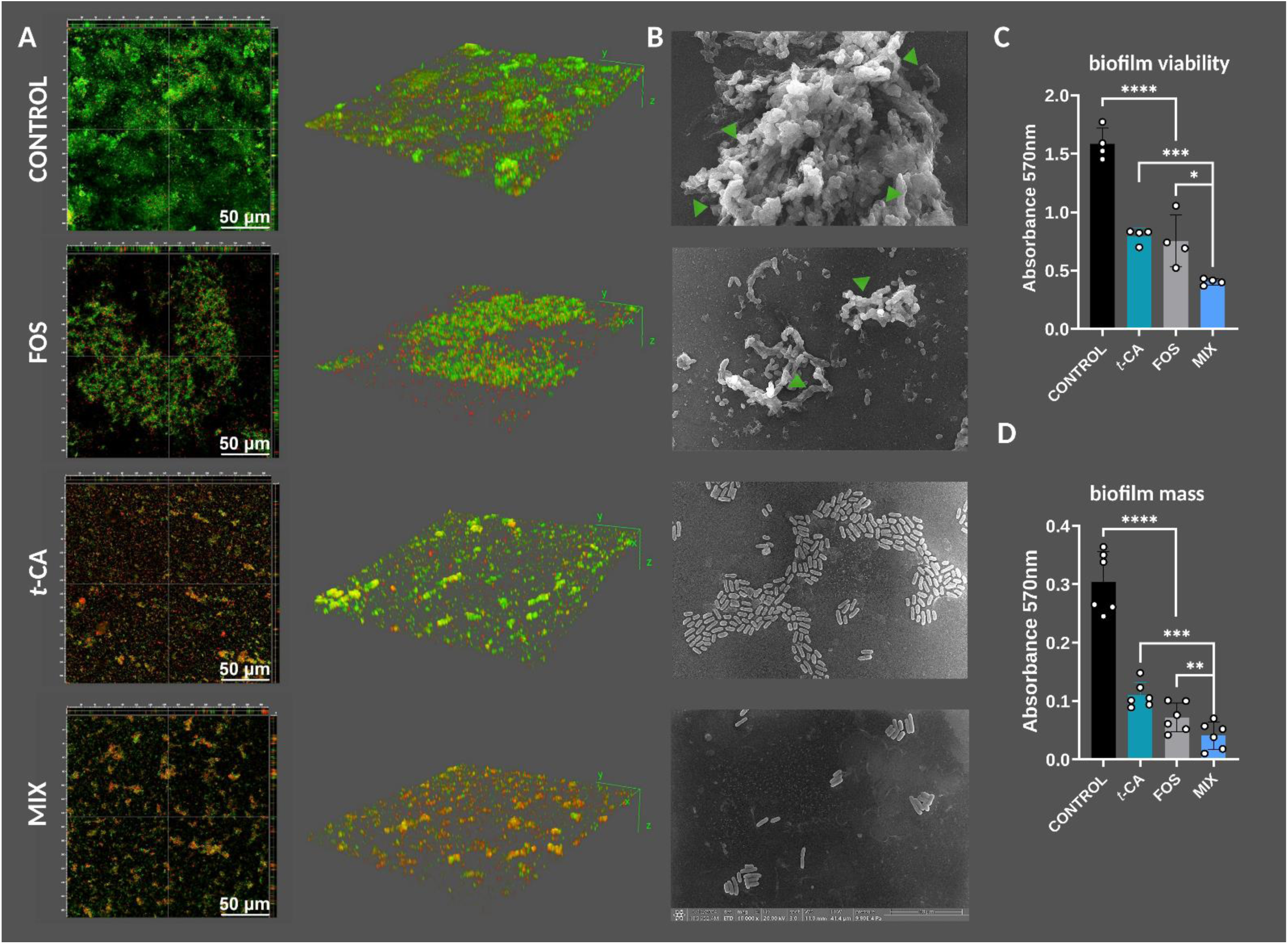
The trans-cinnamaldehyde/fosfomycin combination disrupts three-dimensional biofilm architecture and catheter-associated UPEC biofilms. (A) Representative confocal laser scanning microscopy images and three-dimensional reconstructions of UTI89 biofilms treated with fosfomycin (FOS), trans-cinnamaldehyde (t-CA), or their combination (MIX). Biofilms were stained with SYTO 9 and propidium iodide (PI). SYTO 9-positive cells are shown in green, whereas PI-positive membrane-compromised or dead cells are shown in red. Scale bar: 50 µm. (B) Representative scanning electron microscopy images of catheter-associated UTI89 biofilms after treatment with FOS, t-CA, or MIX. Green arrowheads indicate bacterial aggregates and extracellular biofilm material. (C) Biofilm viability determined by the MTT reduction assay and expressed as absorbance at 570 nm. (D) Total biofilm biomass quantified by crystal violet staining and expressed as absorbance at 570 nm. Data are presented as mean ± SD from independent biological replicates. Statistical significance was assessed as indicated in the figure; *p < 0.05, **p < 0.01, ***p < 0.001, ****p < 0.0001.

### t-CA alters the properties of the cell surface and membrane polarization without inducing ROS

Next, to provide mechanistic insight into the biofilm-associated phenotypes observed above, we investigated whether the reduced viability of biofilm cells was linked to oxidative stress and whether the alterations in extracellular matrix organization were accompanied by changes in cell-surface properties. We therefore assessed intracellular ROS production, cell-surface hydrophobicity, and membrane polarization following exposure to t-CA, FOS, and their combination.

Intracellular ROS production was assessed using DCFDA (Figure 5A). None of the tested concentrations of t-CA, FOS, or their combinations increased fluorescence relative to the untreated control, whereas H₂O₂ produced a pronounced signal. These results indicate that the antibacterial activity of t-CA and the t-CA–FOS combination was not associated with detectable oxidative stress under the tested conditions.

**Figure 5.**
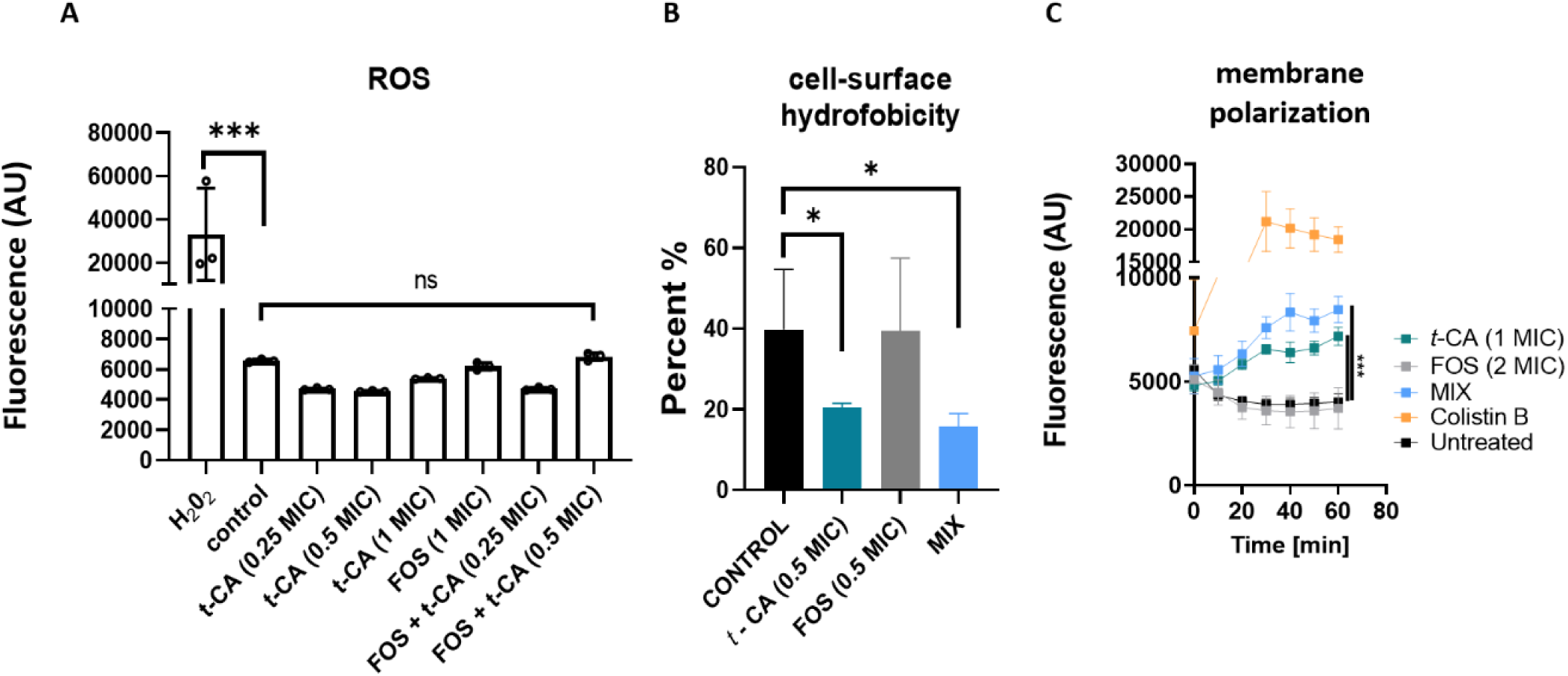
Trans-cinnamaldehyde alters cell-surface properties and membrane polarization without inducing oxidative stress. (A) Intracellular reactive oxygen species (ROS) production in UTI89 cells exposed to trans-cinnamaldehyde (t-CA), fosfomycin (FOS), or their combination, measured using 2′,7′-dichlorofluorescein diacetate (DCFDA). Hydrogen peroxide (H₂O₂) was used as a positive control. (B) Cell-surface hydrophobicity determined using the microbial adhesion to hydrocarbons (MATH) assay. Results were determined spectrophotometrically (A_550_) and presented as a percent of hydrophobicity, tested against untreated control (C) Changes in membrane polarization monitored using the membrane-potential-sensitive fluorescent probe DiSC₃(5). Colistin B was used as a positive control of depolarization. Data are presented as mean ± SD from independent biological replicates. Statistical significance is indicated in the figure; ns, not significant; *p < 0.05, ***p < 0.001.

Cell-surface hydrophobicity was evaluated using the microbial adhesion to hydrocarbons (MATH) assay (Figure 5B). Exposure to t-CA significantly reduced surface hydrophobicity compared with the untreated control, while FOS alone had no detectable effect. A similar reduction was observed for the combination, indicating that the change in surface properties was primarily attributable to t-CA. This finding is consistent with the biofilm-associated alterations observed previously and suggests that t-CA affects bacterial surface features involved in adhesion and biofilm organization.

Membrane polarization was monitored using the potential-sensitive dye DiSC₃(5) (Figure 5C). t-CA induced a gradual increase in fluorescence, consistent with partial membrane depolarization, whereas FOS alone produced only a minor change relative to untreated cells. The combination caused a stronger fluorescence increase than t-CA alone, suggesting an enhanced disturbance of membrane potential. Colistin B, used as a positive control, produced the most pronounced and rapid depolarization.

### t-CA targets the pyruvate node and induces compensatory metabolic alterations

Previous work from our group indicated that t-CA affects central carbon metabolism in *E. coli*, including the upregulation of *poxB*, which encodes pyruvate oxidase (Karczewska *et al*., 2024). Because FOS activity is influenced by the physiological and metabolic state of the cell, including pathways associated with pyruvate utilization, we investigated whether t-CA could interact with enzymes operating at the pyruvate node. Molecular docking was therefore performed against PoxB and three catalytic components of the pyruvate dehydrogenase complex: AceE, AceF, and LpdA (Figure 6.)

**Figure 6.**
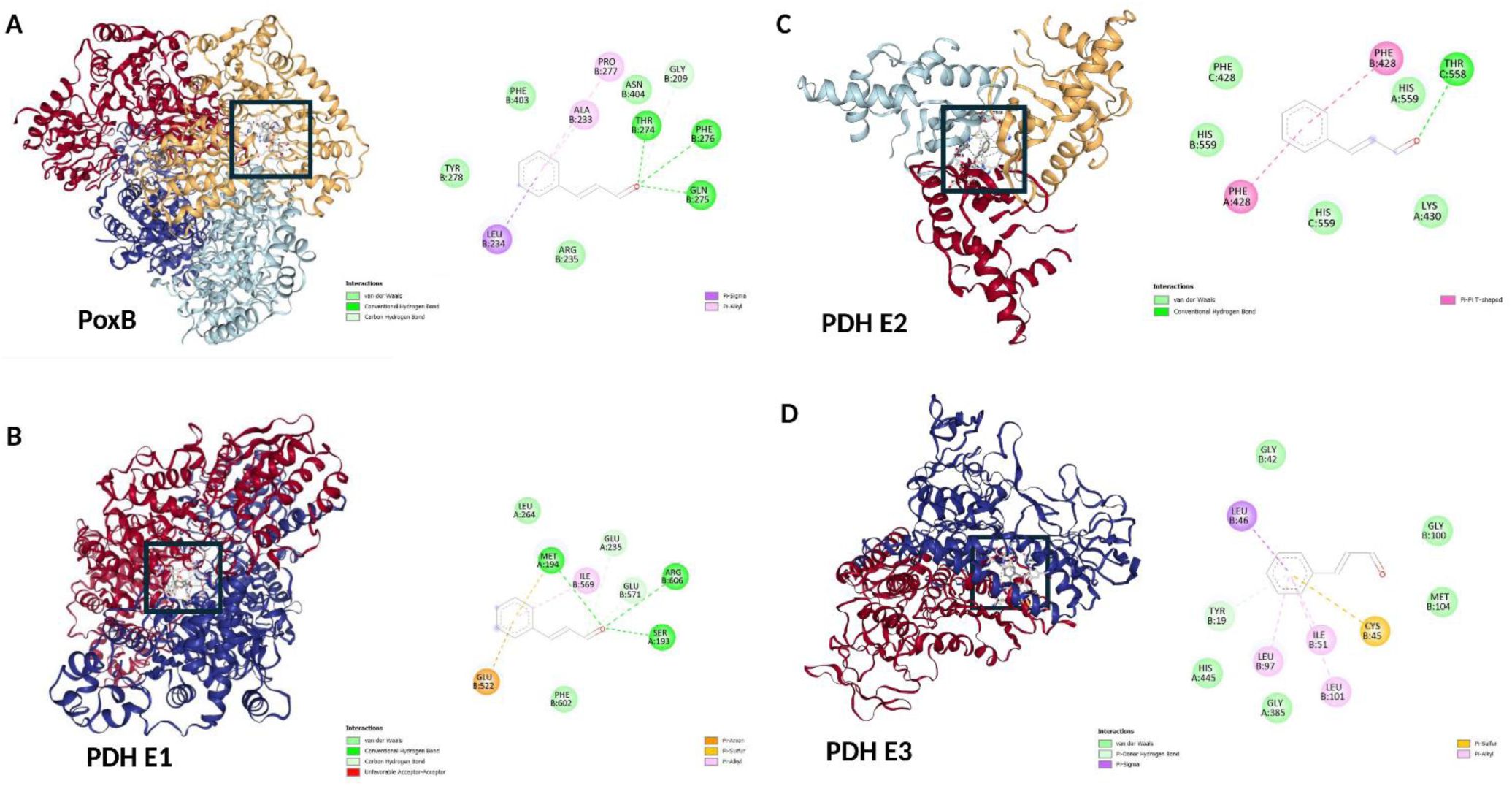
Predicted interactions of trans-cinnamaldehyde with enzymes operating at the pyruvate node. Molecular docking analysis of trans-cinnamaldehyde (t-CA) with enzymes involved in pyruvate metabolism in *E. coli*. Predicted binding poses are shown for (A) pyruvate oxidase (PoxB), (B) the E1 component of the pyruvate dehydrogenase complex (PDH E1; AceE), (C) the E2 component of the pyruvate dehydrogenase complex (PDH E2; AceF), and (D) the E3 component of the pyruvate dehydrogenase complex (PDH E3; LpdA).

Docking predicted energetically favorable poses for all four targets, with calculated binding energies ranging from −5.8 to −6.7 kcal/mol (Table 2). The lowest docking score was obtained for AceF, followed by PoxB, LpdA, and AceE. However, the biological relevance of the predicted interactions differed between the proteins. In AceE, t-CA was positioned within the TPP-binding cavity, adjacent to the catalytic region, suggesting that ligand binding at this site could interfere with cofactor positioning, substrate access, or local conformational dynamics.

**Table 1.** Interaction between t-CA and FOS against laboratory, reference *E. coli*, and clinical UPEC strains. MIC values of t-CA and FOS applied individually and in combination were determined using the checkerboard microdilution assay.

| STRAIN | *Trans*-cinnamaldehyde<br>[mg/l] |  |  | Fosfomycin<br>[mg/l] |  |  | FICI | Interaction |
| --- | --- | --- | --- | --- | --- | --- | --- | --- |
|  | One-component | Combination | Fractional Inhibitory Concentration [FIC] | One-component | Combination | Fractional Inhibitory Concentration [FIC] |  |  |
| MG1655 | 500 | 62.5 | 0.125 | 4 | 1 | 0.25 | 0.375 | synergistic |
| UTI89 | 500 | 16 | 0.03125 | 4 | 1 | 0.25 | 0.28125 | synergistic |
| CFT073 | 500 | 125 | 0.25 | 4 | 1 | 0.25 | 0.5 | synergistic |
| KB077 | 500 | 62.5 | 0.125 | 4 | 1 | 0.25 | 0.375 | synergistic |
| KB075 | 500 | 31 | 0.0625 | 2 | 1 | 0.5 | 0.5625 | additive |
| KB076 | 500 | 125 | 0.25 | 2 | 0.5 | 0.25 | 0.5 | synergistic |
| KB082 | 500 | 31 | 0.0625 | 4 | 1 | 0.25 | 0.3125 | synergistic |
| KB083 | 500 | 16 | 0.03125 | 4 | 2 | 0.5 | 0.53125 | additive |
| KB097 | 500 | 125 | 0.25 | 4 | 0.5 | 0.125 | 0.375 | synergistic |
| KB047 | 500 | 62.5 | 0.125 | 4 | 1 | 0.25 | 0.375 | synergistic |

**Table 2.** Predicted interactions of t-CA with enzymes involved in pyruvate metabolism in *E. coli*. Docking scores and cavity volumes were obtained using CB-Dock2, while ligand–protein contacts were identified with BIOVIA Discovery Studio Visualizer.

| Target | kcal/mol | AAs interactions | Cavity volume (Å<sup>3</sup>) | Contact residues |
| --- | --- | --- | --- | --- |
| PoxB | -6.4 | 6 | 4895 | **Chain A:** THR25 GLY26 ASP27 GLU50 SER73 CYS74 PRO76 GLY77 HIS80 PHE112 GLN113<br>**Chain B:** ASN83 HIS92 GLY209 GLY211 ALA233 LEU234 ARG235 THR251 GLY252 LEU253 ILE254 GLY255 GLY273 THR274 GLN275 PHE276 PRO277 TYR278 ILE291 ASP292 ILE293 ASN294 SER297 GLY310 ASP311 ILE312 VAL380 GLY381 THR384 VAL385 SER402 PHE403 ASN404 GLY406 SER407 MET408 GLY432 ASP433 GLY434 GLY435 MET438 ASN460 VAL462 LEU463 GLY464 PHE465 VAL466 |
| AceE (PDH E1) | -5.8 | 8 | 1776 | **Chain A:** HIS106 SER109 SER112 GLN140 HIS142 TYR177 VAL192 SER193 MET194 GLY229 ASP230 GLY231 GLU232 GLU235 ASN258 ASN260 GLN262 ARG263 LEU264 LYS392 ASP609<br>**Chain B:** ALA520 ASP521 GLU522 ILE569 GLU571 TYR599 PHE602 ARG606 GLU636 HIS640 |
| AceF (PDH E2) | -6.7 | 3 | 2601 | **Chain A:** PHE428 ASP429 LYS430 THR558 HIS559 ALA578 MET579 GLU580 PRO581 VAL582 TRP583 MET593<br>**Chain B:** PHE428 LYS430 THR558 HIS559 ALA578 MET579 GLU580 PRO581 VAL582 TRP583 MET593<br>**Chain C:** PHE428 LYS430 THR558 HIS559 LYS576 SER577 ALA578 MET579 GLU580 PRO581 PRO595 |
| LpdA (PDH E3) | -6.1 | 5 | 1424 | **Chain A:** ALA382 ALA383 SER384 GLY385 ILE388 GLU436 ALA439 LEU440 ILE442 HIS445 GLU460 SER462 ASP465 LEU466 PRO467<br>**Chain B:** PRO16 TYR19 ARG24 GLY42 CYS45 LEU46 ILE51 GLN96 LEU97 GLY100 LEU101 MET104 HIS326 HIS329 VAL330 GLU333 TYR341 PHE342 ASP343 PRO344 |

Although the predicted energy for AceE was less favorable than that obtained for AceF, the location of the binding pose provides a stronger mechanistic rationale for a potential effect on PDH activity. In contrast, the PoxB-binding cavity was substantially larger and accommodated numerous surrounding residues, making the predicted interaction less indicative of a specific inhibitory pose. Thus, the PoxB docking result was considered supportive of possible association but insufficient to infer direct inhibition.

To determine whether these predicted interactions were accompanied by changes in pyruvate metabolism, intracellular pyruvate levels were measured following exposure to subinhibitory t-CA concentrations (Figure 7A). A significant decrease in the intracellular pyruvate pool was observed in cells treated with t-CA alone and with the t-CA/FOS combination. This finding argues against simple pyruvate accumulation and instead indicates altered pyruvate turnover or redistribution of carbon flux. One possible model is that partial perturbation of AceE-dependent PDH activity induces a compensatory metabolic response involving PoxB. Increased PoxB activity would provide an alternative route for direct conversion of pyruvate to acetate and CO₂, potentially lowering the steady-state pyruvate pool while partially bypassing impaired PDH-dependent flux.

**Figure 7.**
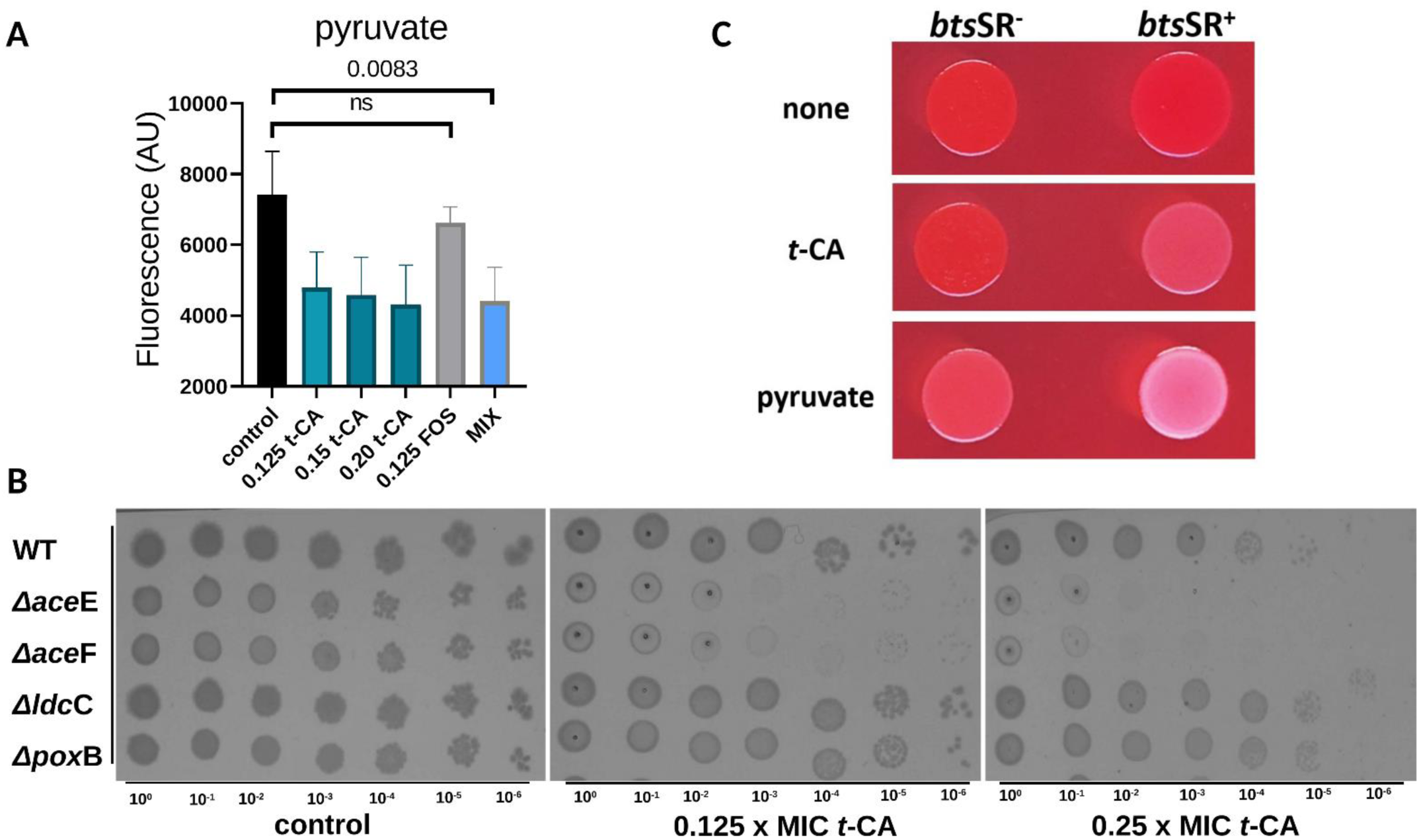
Trans-cinnamaldehyde perturbs pyruvate homeostasis and reveals the contribution of pyruvate metabolism and sensing to the bacterial response. (A) Intracellular pyruvate levels in *E. coli* cells exposed to subinhibitory concentrations of trans-cinnamaldehyde (t-CA), fosfomycin (FOS), or their combination (MIX). Pyruvate levels were measured fluorometrically (Ex:530, Em:585) and expressed as arbitrary fluorescence units (AU). Statistical significance is indicated in the figure; ns, not significant. (B) Serial dilution spot assay of *E. coli* MG1655 wild type (WT) and deletion mutants lacking selected metabolic functions: Δ*aceE*, Δ*aceF*, Δ*ldcC*, and Δ*poxB*. Fresh cultures (OD_600_ = 0.5) were serially diluted from 10⁰ to 10⁻⁶ and spotted onto control plates or plates containing t-CA at 0.125×MIC or 0.25×MIC. (C) Congo red binding phenotype of strains differing in the BtsSR pyruvate-sensing system. Colony phenotypes of *btsSR*⁻ and *btsSR*⁺ backgrounds are shown under untreated conditions, after exposure to t-CA (0.125×MIC), or after supplementation with exogenous pyruvate (60µM). Loss of Congo red binding in the *btsSR*⁺ background indicates that the effect of t-CA and pyruvate on matrix-associated colony phenotype depends on the BtsSR pyruvate-sensing system.

This interpretation is consistent with our previous observation that t-CA induces *poxB* expression without corresponding changes in overflow pathway (Pta-AckA) (Karczewska *et al*., 2024). Such a transcriptional pattern is more compatible with activation of the direct PoxB-dependent route than with classical acetate overflow through the Pta-AckA pathway. Increased conversion of pyruvate to acetate may also explain the previously observed engagement of the lysine decarboxylase-dependent acid stress response. Acetate production and extracellular acidification could activate the Lysine decarboxylation system, which counteracts low-pH stress through proton-consuming lysine decarboxylation. Thus, to further evaluate whether t-CA affects pathways linked to pyruvate metabolism and acid-stress adaptation, spot assays were performed using deletion mutants impaired in metabolic functions selected for the previous test (Figure 7B). Under control conditions, all tested strains retained growth across serial dilutions, although strain-dependent differences in colony density were visible. Exposure to subinhibitory concentrations of t-CA revealed a clear phenotype in mutants defective in PDH components. In particular, disruption of *aceE* and *aceF* markedly impaired growth in the presence of t-CA, indicating that an already compromised PDH complex strongly sensitizes *E. coli* to t-CA exposure (Figures 7B). This phenotype was subsequently confirmed in the UPEC strain UTI89 and its isogenic PDH mutants (Figure S3). This was particularly evident for strains defective in *aceE*, *aceF*, or *lpdA*, supporting the conclusion that PDH-linked pyruvate metabolism contributes to tolerance of t-CA/FOS-mediated stress in a clinically relevant UPEC background.

In contrast, deletion of *ldcC* or *poxB* did not increase susceptibility to t-CA. The growth of these mutants remained broadly comparable to that of the WT strain under the tested conditions. This suggests that LdcC-dependent lysine decarboxylation and PoxB-dependent pyruvate oxidation are unlikely to represent primary cellular targets of t-CA. Rather, these pathways may function as downstream or compensatory responses activated during t-CA-induced metabolic stress.

Additional support for the involvement of pyruvate homeostasis was obtained using strains differing in the BtsSR pyruvate-sensing system (Figure 7C). The contribution of BtsSR to the t-CA-induced phenotype was examined employing *btsSR*-deficient strains. In the presence of a functional BtsSR system, both t-CA (0.125 x MIC) and exogenous pyruvate (60µM) abolished Congo red binding, consistent with reduced production of matrix-associated components. This response was not observed in the *btsSR*-deficient background, which retained the Congo red-positive phenotype under both treatment conditions. Thus, the effects of t-CA and pyruvate on biofilm matrix production were dependent on BtsSR-mediated pyruvate sensing.

### The t-CA/FOS combination attenuates the development of fosfomycin resistance

The emergence of antimicrobial resistance is a major concern in the treatment of recurrent UTIs, where repeated antibiotic exposure may promote the selection of less susceptible bacterial populations (Figure 8). Serial exposure to FOS alone resulted in a rapid and progressive increase in MIC, reaching approximately 300-fold above the initial value by day 5. The addition of t-CA delayed this increase in a concentration-dependent manner, with 0.25×MIC t-CA providing the strongest suppression of development of FOS resistance. In contrast, repeated exposure to t-CA alone did not result in a detectable increase in MIC or reduced susceptibility. Thus, the finding that, t-CA limited the development of reduced susceptibility to FOS without itself promoting resistance during serial passage was both unexpected and intriguing. We next asked whether t-CA could improve the activity of FOS against FOS-resistant UPEC strains with different resistance backgrounds (Figure 8B). The combination showed no relevant interaction in strains carrying FosA-type metalloenzymes, including R82 (*fosA3*) and R4880 (*fosA4*), indicating that t-CA did not restore FOS activity when resistance was mediated by enzymatic drug inactivation. In contrast, the FOS-resistant strain N-279, which did not carry a detectable FosA-type determinant, responded positively to the combination. In this strain, the presence of t-CA reduced the effective FOS concentration fourfold, from an initial MIC of 2 g/L to 0.5 g/L.

**Figure 8.**
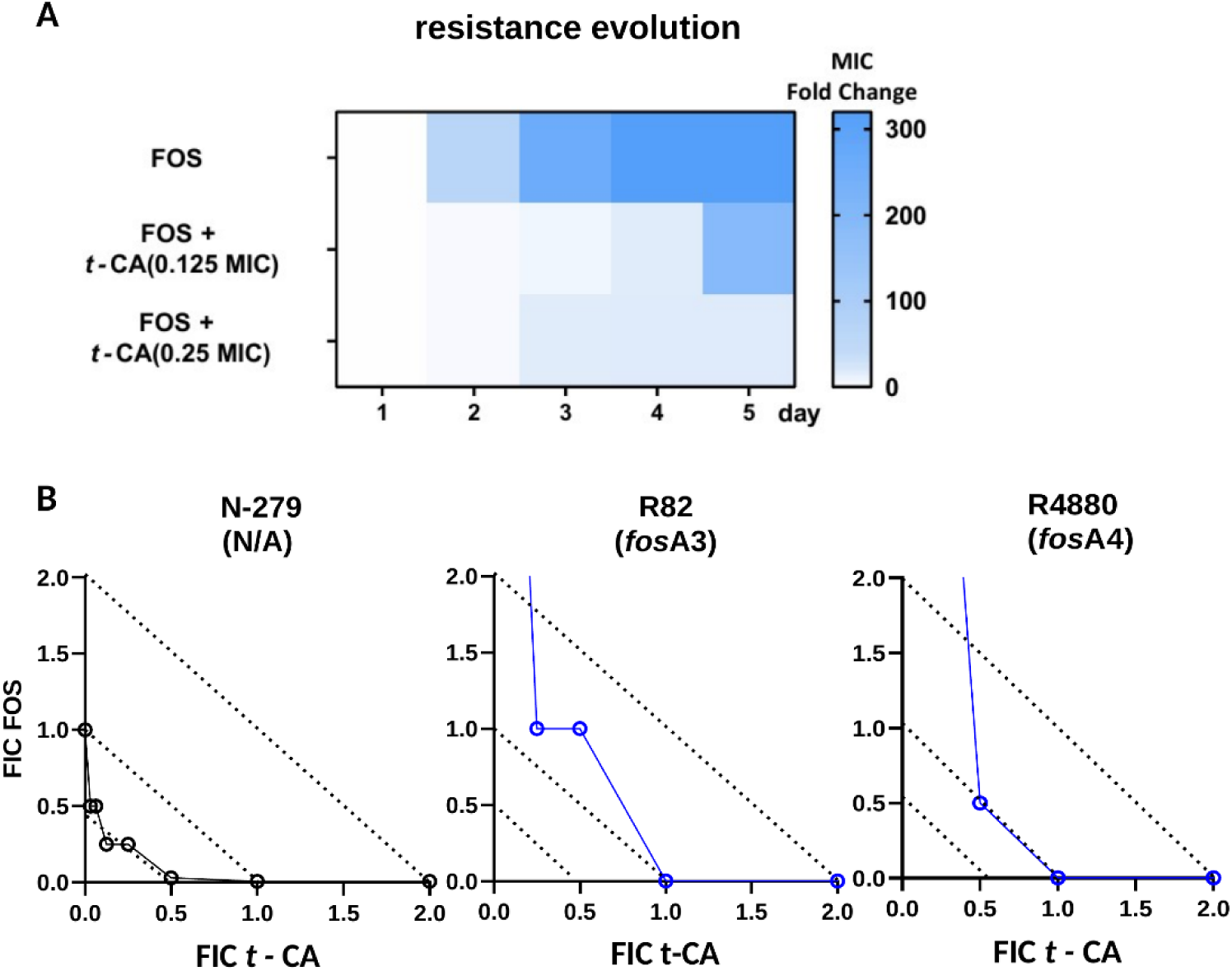
Trans-cinnamaldehyde limits the development of reduced fosfomycin susceptibility and potentiates fosfomycin activity in isolates without FosA-mediated resistance. (A) Experimental evolution of fosfomycin (FOS) susceptibility during serial exposure. *E. coli* cultures were passaged for 5 consecutive days in the presence of FOS alone or FOS combined with trans-cinnamaldehyde (t-CA) at 0.125×MIC or 0.25×MIC. (B) Isobolograms show the interaction between FOS and t-CA in FOS-resistant clinical isolates with different resistance backgrounds (N-279, which lacked detectable *fosA1–fosA6* determinants, R82 and R4880, carrying the FosA-type metalloenzyme genes *fosA3* and *fosA4*, respectively).

### The t-CA/FOS combination treatment improves survival in the *G. mellonella* infection model

The protective effect of t-CA and FOS was further evaluated in a *G. mellonella* model of UTI89 infection as presented on schematic pipeline (Figure 9A). Infection (5 x 10^6^ cfu/larvae) followed by PBS treatment resulted in progressive larval mortality, whereas untreated non-infected controls remained fully viable throughout the experiment (Figure 9B). FOS monotherapy improved survival in a dose-dependent manner, although the effect remained partial. In contrast, t-CA alone did not significantly improve survival under the tested conditions. The strongest protective effect was observed for the combined treatment. All tested t-CA/FOS combinations maintained survival at, or close to, 100% during the observation period, with the lowest-dose combination showing only a limited decline at the final time point. These results indicate that t-CA markedly enhances the *in vivo* efficacy of FOS against virulent UPEC pathogen UTI89 infection, even though presents limited protective activity when administered alone.

**Figure 9.**
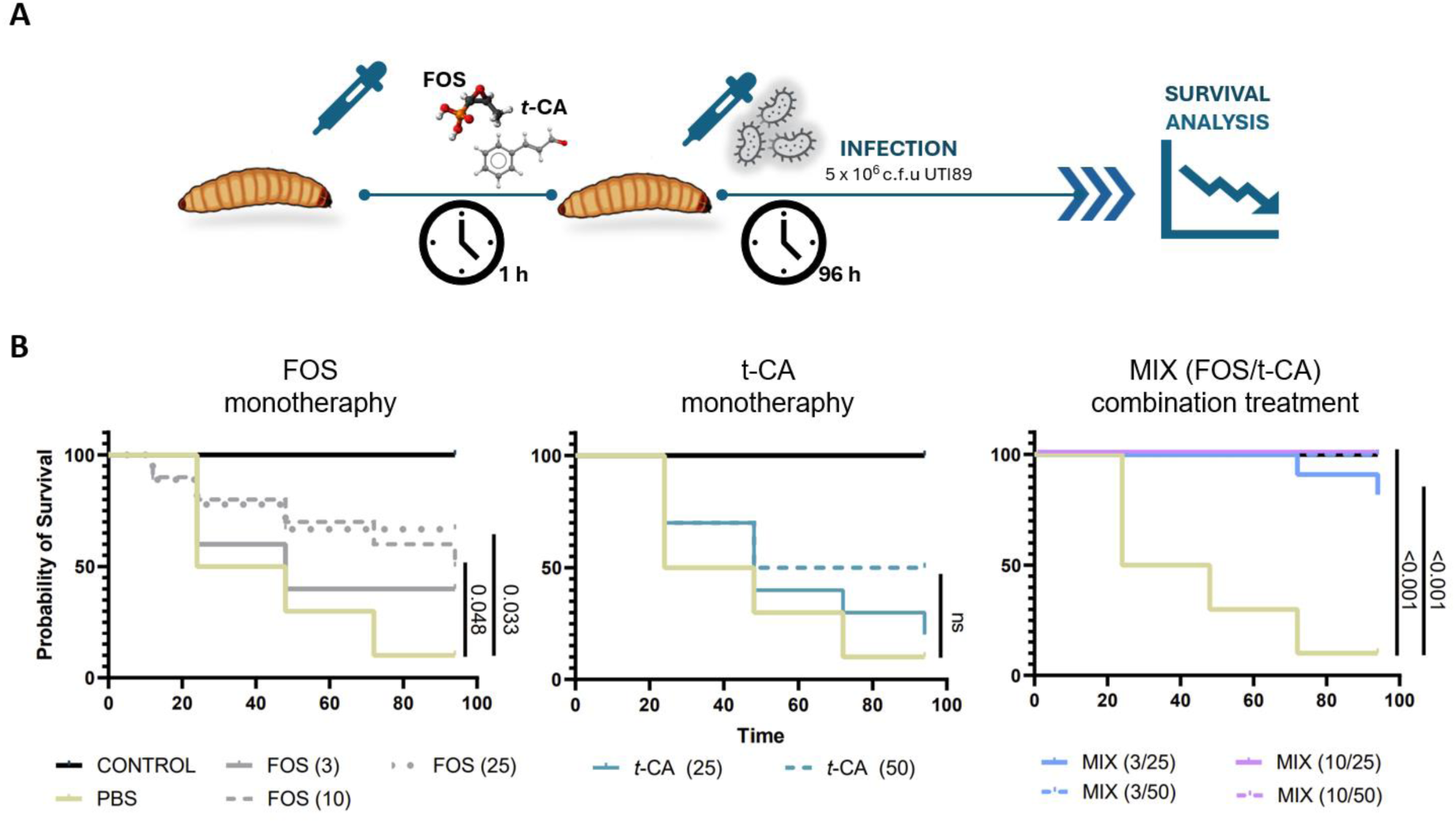
The trans-cinnamaldehyde/fosfomycin combination improves survival in a *Galleria mellonella* model of UTI89 infection. (A) Schematic representation of the experimental workflow. *G. mellonella* larvae were pre-treated with fosfomycin (FOS), trans-cinnamaldehyde (t-CA), or their combination. After 1 h, larvae were infected with the uropathogenic *E. coli* strain UTI89 at 5 × 10⁶ colony-forming units (CFU) per larva. Survival was monitored for 96 h. (B) Kaplan-Meier survival curves of infected larvae treated with FOS, t-CA, or the FOS/t-CA combination. PBS-treated infected larvae and non-infected untreated larvae were included as controls. All doses used in experiment are expressed as mg/kg body weight. Statistical significance was assessed using the log-rank test; ns, not significant.

## 4. Discussion

Recurrent and difficult-to-treat UTIs are strongly associated with UPEC biofilm formation, particularly in hypervirulent strains, where biofilms promote persistence, antimicrobial tolerance, and relapse (Flores-Mireles *et al*., 2015). This problem is especially relevant in complicated settings, including diabetes and urinary tract instrumentation. Accordingly, FOS remains an attractive therapeutic option because of its broad activity against uropathogens, high urinary exposure, low side-effect for patients and clinical usefulness in complicated cases. Recent data from Ye et al. showed that fosfomycin trometamol reduced biofilm-forming bacterial colonization on double-J stents in diabetic patients compared with levofloxacin (Ye *et al*., 2026).

Our results show that t-CA potentiates the antibiofilm activity of FOS. While FOS reduced biofilm growth and viability, t-CA additionally depleted ECM and disrupted biofilm architecture, consistent with our recent transcriptomic data showing downregulation of genes involved in fimbriae and ECM production (Karczewska *et al*., 2026). This simple mechanistic outcome is relevant because biofilm penetration is a major determinant of antimicrobial efficacy. Dzib-Baak et al. demonstrated that FOS activity against UPEC biofilms depends on the biofilm-producing phenotype, with stronger biofilm producers requiring higher FOS concentrations for biofilm degradation (Dzib-Baak *et al*., 2022). Thus, t-CA-mediated weakening of the biofilm matrix may improve FOS efficacy by reducing the structural barrier protecting biofilm-associated cells. Conversely, inadequate exposure to FOS may favor biofilm persistence or even enhance biofilm formation under sub-inhibitory conditions, as shown for *Staphylococcus aureus* through a SarA-dependent mechanism (Zeng *et al*., 2025). This further supports the rationale for combining FOS with an agent that simultaneously reduces biofilm matrix integrity.

The envelope-associated effects observed in our study are consistent with this interpretation. The t-CA–FOS combination affected cell-surface hydrophobicity and membrane potential, but membrane depolarization occurred more slowly than in the colistin-treated control. This suggests that the combination does not act primarily as a rapidly membrane-disrupting, detergent-like treatment. Instead, the slower kinetics may reflect t-CA-driven remodeling of surface-associated structures, local loosening of the cell envelope, and gradual leakage of membrane-potential-sensitive dye. Such an effect would be compatible with reduced ECM abundance, altered biofilm cohesion, and increased physiological stress in biofilm-associated cells.

Together, our data support a model in which t-CA perturbs the pyruvate node rather than acting solely as a membrane-active compound. Docking placed t-CA in the TPP-associated cavity of AceE, close to the catalytic region, while metabolite analysis showed reduced intracellular pyruvate after t-CA treatment. One possible explanation for this apparently counterintuitive decrease is compensatory rerouting of pyruvate through PoxB, whose expression is increased upon t-CA exposure (Chang *et al*., 1994; Karczewska *et al*., 2024, 2026). In such a model, PoxB-dependent conversion of pyruvate to acetate and CO₂ could partially bypass the canonical PDH-dependent route to acetyl-CoA and lower the steady-state pyruvate pool (Ogasawara et al., 2007; Sauer & Eikmanns, 2005). Such rerouting may lower the steady-state pyruvate pool, increase acetate-associated acid stress, and engage lysine decarboxylase-dependent pH homeostasis (Moreau, 2007). This interpretation is further supported by recent work in *Streptococcus mutans*, where t-CA was shown to interfere with carbohydrate metabolism, glycolysis, pyruvate metabolism, and the TCA cycle (Zhang *et al*., 2024). In that study, proteomic analysis, metabolic assays, qRT-PCR, and molecular docking collectively identified PDH as a potential target of t-CA, and t-CA exposure reduced sugar utilization, LDH activity, and ATP production. The involvement of pyruvate sensing is further supported by our observation that both t-CA and exogenous pyruvate abolished Congo red binding only in the BtsSR-proficient background (Vilhena *et al*., 2018). Because FOS activity depends on bacterial physiology and drug uptake, including outer-membrane porins and metabolic state, this t-CA-driven metabolic imbalance may reduce the capacity of *E. coli* to tolerate MurA inhibition and thereby potentiate FOS activity (Bianchi *et al*., 2024). This interpretation is consistent with recent work by Verhülsdonk et al., who showed that the metabolic state of *E. coli* modulates FOS efficacy and promotes resistance evolution. Importantly, their study identified *btsT*, encoding the pyruvate transporter controlled by the BtsSR system, among the upregulated genes associated with post-FOS regrowth (Verhulsdonk *et al*., 2026). In this context, our observation that both t-CA and exogenous pyruvate abolish Congo red binding in a BtsSR-dependent manner suggests that t-CA intersects with the same pyruvate-sensing axis that can influence recovery after FOS exposure. Thus, t-CA-driven perturbation of pyruvate homeostasis may not only potentiate the immediate antibacterial effect of FOS, but also interfere with metabolic states permissive for regrowth and resistance evolution. This conclusion is further supported by the response of FOS-resistant isolates with different resistance backgrounds (Mueller *et al*., 2019; Nordmann *et al*., 2019). Strains carrying genes encoding FosA-type metalloenzymes, including FosA3 and FosA4, did not show detectable re-sensitization to FOS in the presence of t-CA. In contrast, isolate N-279, whose FOS resistance was not associated with enzymatic drug inactivation, displayed restored susceptibility under combined treatment. This distinction suggests that t-CA does not overcome resistance mediated by direct FosA-dependent inactivation of FOS. Rather, its potentiating effect appears to be most relevant when reduced FOS susceptibility is linked to metabolic or physiological adaptation, consistent with the proposed role of t-CA in perturbing pyruvate homeostasis and FOS-responsive recovery pathways.

The present data also suggest that t-CA does not promote FOS resistance under the tested conditions. Although t-CA was previously shown to induce RelA-dependent (p)ppGpp accumulation, this response was not accompanied by activation of major resistance-associated genes, including *marA*, *acrD*, *ampC*, or efflux-related determinants (Karczewska *et al*., 2024, 2026). It is intriguing in the light of data showing increased antibiotic tolerance upon induction of the stringent response (Spira & Ospino, 2020), and this phenomenon was reported for ciprofloxacin, chloramphenicol, rifampicin, mecillinam, ampicillin, and trimethoprim. For FOS, repeated exposure to t-CA alone did not increase MIC, whereas the combination of t-CA with FOS limited the progressive MIC elevation observed during exposure to FOS alone. Thus, t-CA-induced stress signalling does not appear to translate into a pro-resistance phenotype. This interpretation is consistent with previous studies showing that t-CA can enhance antibacterial activity without promoting stable resistance, including activity against biofilms and multidrug-resistant bacteria (Qiu et al., 2026; Usai & Di Sotto, 2023). Moreover, t-CA was recently shown to target the staphylococcal accessory regulator SarA, thereby enhancing β-lactam activity against methicillin-resistant *Staphylococcus aureus* (Li *et al*., 2024). Prolonged exposure of t-CA, resulting in parallelly elevated (p)ppGpp level would be considered as a stabilizing factor for DNA repair, thus decreasing the level of spontaneous mutations which could potentially lead to antibiotic resistance (Sivapragasam, 2025; Weaver *et al*., 2023). In our model, the effects of t-CA on pyruvate homeostasis, biofilm matrix production, and metabolic recovery may similarly constrain adaptive states required for FOS tolerance and regrowth. However, this conclusion remains provisional and requires further validation, including analysis of evolved populations, FOS transporter expression, and (p)ppGpp dynamics during combined treatment.

The potential clinical relevance of this interaction is supported by the use of two complementary infection-associated models. First, the catheter biofilm model showed that the t-CA/FOS combination was effective against surface-associated UPEC communities, which are directly relevant to catheter-associated and device-associated UTIs. Second, the improved survival of infected *G. mellonella* larvae indicates that the enhanced activity of the combination is not limited to in vitro biofilm assays but is also observed in a living host model. This is relevant because *G. mellonella* is an established surrogate infection model widely used for screening bacterial virulence and evaluating novel antimicrobial strategies, including in UTI-related studies (Chitolina *et al*., 2025; Karczewska *et al*., 2023; Pereira *et al*., 2020). Nevertheless, an important limitation of this study is that the insect model does not reproduce the anatomical, immunological, and pharmacokinetic complexity of mammalian UTIs. However, even if the direct translation of the results obtained in the *G. mellonella* is not possible, also because in this simplified infection model biofilm formation is not usually a key player, it is apparent that the FOS/t-CA combination effect involves decreased virulence even before pathogens can form more complex structure. Therefore, the improved survival of infected larvae should be interpreted as proof-of-concept evidence supporting the in vivo relevance of the t-CA/FOS interaction, rather than as direct evidence of therapeutic efficacy in the urinary tract. Further validation in mammalian UTI models will be required to determine whether t-CA can enhance the therapeutic potential of FOS against biofilm-associated UPEC infections.

## 5. Conclusions

This study demonstrates that t-CA acts as a phytochemical adjuvant that enhances FOS activity against biofilm-forming *E. coli*, including UPEC strains. The main novelty of this work is the identification of a dual mode of potentiation: t-CA weakens the biofilm matrix while simultaneously perturbing bacterial metabolic adaptation at the pyruvate node. This combined antibiofilm and metabolic effect increased FOS efficacy, improved the outcome in an in vivo infection model, and, importantly, limited the emergence of reduced FOS susceptibility during repeated exposure. Our findings support a model in which plant-derived metabolites can enhance antibiotic activity not only by direct antibacterial or antibiofilm effects, but also by restricting bacterial physiological plasticity and adaptive recovery. By targeting both biofilm-associated protection and metabolic states permissive for FOS tolerance, t-CA may help reduce the resistance-developing potential of FOS-based treatment strategies. Nonetheless, further validation in mammalian UTI models and evolved bacterial populations will be required to confirm the therapeutic relevance and resistance-limiting potential of this approach.

## Acknowledgments

We are grateful to prof. Kirsten Jung for providing *bts*SR and *bts*T *E. coli* mutants used in this work, prof. Patrice Nordmann for providing fosfomycin resistant UPEC isolates and Dorota Łuszczek for her technical support with SEM imagining.

## 6. Authors Contribution

**Monika Karczewska**: Conceptualization, Investigation, Methodology, Data Curation, Visualization, Writing – Original Draft. **Patryk Strzelecki**: Investigation. **Monika Maciąg-Dorszyńska**: Investigation, Methodology, Data Curation. **Małgorzata Kapusta**: Investigation, Methodology, Data Curation. **Agnieszka Pyrczak-Felczykowska:** Investigation, Methodology **Agnieszka Szalewska-Pałasz**: Methodology, Validation, Supervision, Writing – Review & Editing. **Dariusz Nowicki**: Conceptualization, Investigation, Methodology, Validation, Visualization, Funding Acquisition, Supervision, Writing – Original Draft, Writing – Review & Editing.

## 7. Declaration statement

We declare no conflicts of interest related to this study.

## 8. Data availability

All original data generated during the study are included in the article in the form of figures and tables. For any further inquiries, please contact the corresponding author.

## Abbreviations

AMR: antimicrobial resistance
AU: arbitrary units
CFU: colony-forming units
CLSM: confocal laser scanning microscopy
CV: crystal violet
DCFDA: 2′,7′-dichlorofluorescein diacetate
DMSO: dimethyl sulfoxide
ECM: extracellular matrix
FIC: fractional inhibitory concentration
FICI: fractional inhibitory concentration index
FOS: fosfomycin
GSH: reduced glutathione
MATH: microbial adhesion to hydrocarbons
MBC: minimum bactericidal concentration
MDR: multidrug-resistant
MIC: minimum inhibitory concentration
MTT: 3-(4,5-dimethylthiazol-2-yl)-2,5-diphenyltetrazolium bromide
OD: optical density
PBS: phosphate-buffered saline
PDH: pyruvate dehydrogenase complex
PEP: phosphoenolpyruvate
PI: propidium iodide
ROS: reactive oxygen species
SEM: scanning electron microscopy
t-CA: trans-cinnamaldehyde
TPP: thiamine pyrophosphate
UPEC: uropathogenic *Escherichia coli*
UTIs: urinary tract infections
WT: wild type.

## References

1. Bianchi, M., Winterhalter, M., Harbig, T.A., Horompoli, D., Ghai, I., Nieselt, K., Brotz-Oesterhelt, H., et al. (2024), “Fosfomycin Uptake in Escherichia coli Is Mediated by the Outer-Membrane Porins OmpF, OmpC, and LamB”, ACS Infect Dis, Vol. 10 No. 1, pp. 127–137, doi: 10.1021/acsinfecdis.3c00367.

2. Borowicz, M., Krzyzanowska, D.M. and Jafra, S. (2023), “Crystal violet-based assay for the assessment of bacterial biofilm formation in medical tubing”, J Microbiol Methods, Vol. 204, p. 106656, doi: 10.1016/j.mimet.2022.106656.

3. Budziaszek, J., Pilarczyk-Zurek, M., Dobosz, E., Kozinska, A., Nowicki, D., Obszanska, K., Szalewska-Palasz, A., et al. (2023), “Studies of Streptococcus anginosus virulence in Dictyostelium discoideum and Galleria mellonella models”, Infect Immun, Vol. 91 No. 5, p. e0001623, doi: 10.1128/iai.00016-23.

4. Buttress, J.A., Halte, M., Te Winkel, J.D., Erhardt, M., Popp, P.F. and Strahl, H. (2022), “A guide for membrane potential measurements in Gram-negative bacteria using voltage-sensitive dyes”, Microbiology, Vol. 168 No. 9, doi: 10.1099/mic.0.001227.

5. Chang, Y.Y., Wang, A.Y. and Cronan Jr., J.E. (1994), “Expression of Escherichia coli pyruvate oxidase (PoxB) depends on the sigma factor encoded by the rpoS(katF) gene”, Mol Microbiol, Vol. 11 No. 6, pp. 1019–1028, doi: 10.1111/j.1365-2958.1994.tb00380.x.

6. Chitolina, G.Z., Furian, T.Q., Granados, O.F.O., Borges, K.A., Weber, T.B., Goncalves, I.B., Mollerke, R., et al. (2025), “Determining the pathogenicity of uropathogenic Escherichia coli strains in the Galleria mellonella larvae model”, Altern Lab Anim, Vol. 53 No. 4, pp. 203–214, doi: 10.1177/02611929251349791.

7. CLSI, C. (2018), “Methods for dilution antimicrobial susceptibility tests for bacteria that grow aerobically”, Clinical and Laboratory Standards Institute Wayne, PA.

8. Dzib-Baak, H.E., Uc-Cachon, A.H., Dzul-Beh, A.J., Rosado-Manzano, R.F., Gracida-Osorno, C. and Molina-Salinas, G.M. (2022), “Efficacy of Fosfomycin against Planktonic and Biofilm-Associated MDR Uropathogenic Escherichia coli Clinical Isolates”, Trop Med Infect Dis, Vol. 7 No. 9, doi: 10.3390/tropicalmed7090235.

9. Falagas, M.E., Kastoris, A.C., Kapaskelis, A.M. and Karageorgopoulos, D.E. (2010), “Fosfomycin for the treatment of multidrug-resistant, including extended-spectrum beta-lactamase producing, Enterobacteriaceae infections: a systematic review”, Lancet Infect Dis, Vol. 10 No. 1, pp. 43–50, doi: 10.1016/S1473-3099(09)70325-1.

10. Flores-Mireles, A.L., Walker, J.N., Caparon, M. and Hultgren, S.J. (2015), “Urinary tract infections: epidemiology, mechanisms of infection and treatment options”, Nat Rev Microbiol, Vol. 13 No. 5, pp. 269–284, doi: 10.1038/nrmicro3432.

11. Gil-Gil, T. and Martinez, J.L. (2022), “Glucose-6-phosphate Reduces Fosfomycin Activity Against Stenotrophomonas maltophilia”, Front Microbiol, Vol. 13, p. 863635, doi: 10.3389/fmicb.2022.863635.

12. Gil-Gil, T. and Martinez, J.L. (2024), “Role of the phosphotransferase system in the transport of fosfomycin in Escherichia coli”, Int J Antimicrob Agents, Vol. 63 No. 1, p. 107027, doi: 10.1016/j.ijantimicag.2023.107027.

13. Groleau, M.C., Houle, S., Quevedo, A.C., McKay, G., Nguyen, D., Dozois, C.M., Tufenkji, N., et al. (2026), “Cranberry juice potentiates sensitivity of uropathogenic Escherichia coli (UPEC) strains to fosfomycin and decreases occurrence of spontaneous resistance”, Appl Environ Microbiol, Vol. 92 No. 5, p. e0252125, doi: 10.1128/aem.02521-25.

14. Iskandar, K., Rizk, R., Matta, R., Husni-Samaha, R., Sacre, H., Bouraad, E., Dirani, N., et al. (2021), “Economic Burden of Urinary Tract Infections From Antibiotic-Resistant Escherichia coli Among Hospitalized Adult Patients in Lebanon: A Prospective Cohort Study”, Value Health Reg Issues, Vol. 24, pp. 38–46, doi: 10.1016/j.vhri.2020.01.006.

15. Karczewska, M., Strzelecki, P., Bogucka, K., Potrykus, K., Szalewska-Palasz, A. and Nowicki, D. (2023), “Increased Levels of (p)ppGpp Correlate with Virulence and Biofilm Formation, but Not with Growth, in Strains of Uropathogenic Escherichia coli”, Int J Mol Sci, Vol. 24 No. 4, doi: 10.3390/ijms24043315.

16. Karczewska, M., Strzelecki, P., Pyrczak-Felczykowska, A., Zima, K., Szalewska-Palasz, A. and Nowicki, D. (2026), “Trans-cinnamaldehyde triggers stringent response-mediated virulence attenuation in pathogenic Escherichia coli”, Biomed Pharmacother, Vol. 198, p. 119306, doi: 10.1016/j.biopha.2026.119306.

17. Karczewska, M., Wang, A.Y., Narajczyk, M., Slominski, B., Szalewska-Palasz, A. and Nowicki, D. (2024), “Antibacterial activity of t-cinnamaldehyde: An approach to its mechanistic principle towards enterohemorrhagic Escherichia coli (EHEC)”, Phytomedicine, Vol. 132, p. 155845, doi: 10.1016/j.phymed.2024.155845.

18. Klepser, M.E., Ernst, E.J., Lewis, R.E., Ernst, M.E. and Pfaller, M.A. (1998), “Influence of test conditions on antifungal time-kill curve results: proposal for standardized methods”, Antimicrob Agents Chemother, Vol. 42 No. 5, pp. 1207–1212, doi: 10.1128/AAC.42.5.1207.

19. Konieczny, M., Rhein, P., Czaczyk, K., Bialas, W. and Juzwa, W. (2021), “Imaging Flow Cytometry to Study Biofilm-Associated Microbial Aggregates”, Molecules, Vol. 26 No. 23, doi: 10.3390/molecules26237096.

20. Li, J., Lu, T., Chu, Y., Zhang, Y., Zhang, J., Fu, W., Sun, J., et al. (2024), “Cinnamaldehyde targets SarA to enhance beta-lactam antibiotic activity against methicillin-resistant Staphylococcus aureus”, MLife, Vol. 3 No. 2, pp. 291–306, doi: 10.1002/mlf2.12121.

21. Liu, Y., Yang, X., Gan, J., Chen, S., Xiao, Z.X. and Cao, Y. (2022), “CB-Dock2: improved protein-ligand blind docking by integrating cavity detection, docking and homologous template fitting”, Nucleic Acids Res, Vol. 50 No. W1, pp. W159–W164, doi: 10.1093/nar/gkac394.

22. Mancuso, G., Midiri, A., Gerace, E., Marra, M., Zummo, S. and Biondo, C. (2023), “Urinary Tract Infections: The Current Scenario and Future Prospects”, Pathogens, Vol. 12 No. 4, doi: 10.3390/pathogens12040623.

23. Mattioni Marchetti, V., Hrabak, J. and Bitar, I. (2023), “Fosfomycin resistance mechanisms in Enterobacterales: an increasing threat”, Front Cell Infect Microbiol, Vol. 13, p. 1178547, doi: 10.3389/fcimb.2023.1178547.

24. Moreau, P.L. (2007), “The lysine decarboxylase CadA protects Escherichia coli starved of phosphate against fermentation acids”, J Bacteriol, Vol. 189 No. 6, pp. 2249–2261, doi: 10.1128/JB.01306-06.

25. Mueller, L., Cimen, C., Poirel, L., Descombes, M.C. and Nordmann, P. (2019), “Prevalence of fosfomycin resistance among ESBL-producing Escherichia coli isolates in the community, Switzerland”, Eur J Clin Microbiol Infect Dis, Vol. 38 No. 5, pp. 945–949, doi: 10.1007/s10096-019-03531-0.

26. Nordmann, P., Poirel, L. and Mueller, L. (2019), “Rapid Detection of Fosfomycin Resistance in Escherichia coli”, J Clin Microbiol, Vol. 57 No. 1, doi: 10.1128/JCM.01531-18.

27. Ogasawara, H., Ishida, Y., Yamada, K., Yamamoto, K. and Ishihama, A. (2007), “PdhR (pyruvate dehydrogenase complex regulator) controls the respiratory electron transport system in Escherichia coli”, J Bacteriol, Vol. 189 No. 15, pp. 5534–5541, doi: 10.1128/JB.00229-07.

28. Pembrey, R.S., Marshall, K.C. and Schneider, R.P. (1999), “Cell surface analysis techniques: What do cell preparation protocols do to cell surface properties?”, Appl Environ Microbiol, Vol. 65 No. 7, pp. 2877–2894, doi: 10.1128/AEM.65.7.2877-2894.1999.

29. Peng, B., Li, H. and Peng, X.X. (2025), “Metabolic state-driven nutrient-based approach to combat bacterial antibiotic resistance”, NPJ Antimicrob Resist, Vol. 3 No. 1, p. 24, doi: 10.1038/s44259-025-00092-5.

30. Pereira, M.F., Rossi, C.C., da Silva, G.C., Rosa, J.N. and Bazzolli, D.M.S. (2020), “Galleria mellonella as an infection model: an in-depth look at why it works and practical considerations for successful application”, Pathog Dis, Vol. 78 No. 8, doi: 10.1093/femspd/ftaa056.

31. Qiu, X., Wang, B. and Wang, Y. (2026), “Cinnamaldehyde attenuates intergeneric horizontal transfer of antibiotic resistance genes by disrupting quorum sensing”, ENGINEERING Environment, Vol. 20 No. 6, p. 85, doi: 10.1007/s11783-026-2185-x.

32. Rahman, M.M., Hossain, M.M.K., Rubaya, R., Halder, J., Karim, M.E., Bhuiya, A.A., Khatun, A., et al. (2022), “Association of Antibiotic Resistance Traits in Uropathogenic Escherichia coli (UPEC) Isolates”, Can J Infect Dis Med Microbiol, Vol. 2022, p. 4251486, doi: 10.1155/2022/4251486.

33. Sauer, U. and Eikmanns, B.J. (2005), “The PEP-pyruvate-oxaloacetate node as the switch point for carbon flux distribution in bacteria”, FEMS Microbiol Rev, Vol. 29 No. 4, pp. 765–794, doi: 10.1016/j.femsre.2004.11.002.

34. Sivapragasam, S. (2025), “The Emerging Role of (p)ppGpp in DNA Repair and Associated Bacterial Survival against Fluoroquinolones”, Gene Expr, Vol. 24 No. 2, pp. 130–138, doi: 10.14218/ge.2024.00033.

35. Spira, B. and Ospino, K. (2020), “Diversity in E. coli (p)ppGpp Levels and Its Consequences”, Front Microbiol, Vol. 11, p. 1759, doi: 10.3389/fmicb.2020.01759.

36. Strzelecki, P., Karczewska, M., Szalewska-Palasz, A. and Nowicki, D. (2025), “Phytochemicals Controlling Enterohemorrhagic Escherichia coli (EHEC) Virulence-Current Knowledge of Their Mechanisms of Action”, Int J Mol Sci, Vol. 26 No. 1, doi: 10.3390/ijms26010381.

37. Usai, F. and Di Sotto, A. (2023), “trans-Cinnamaldehyde as a Novel Candidate to Overcome Bacterial Resistance: An Overview of In Vitro Studies”, Antibiotics, Vol. 12 No. 2, doi: 10.3390/antibiotics12020254.

38. Verhulsdonk, A., Stadelmann, A., Smollich, F., Rapp, J., Straub, D. and Link, H. (2026), “The Metabolic State of E. coli Influences Fosfomycin Efficacy and Promotes Resistance Evolution”, ACS Infect Dis, Vol. 12 No. 3, pp. 1155–1164, doi: 10.1021/acsinfecdis.5c01013.

39. Vilhena, C., Kaganovitch, E., Shin, J.Y., Grunberger, A., Behr, S., Kristoficova, I., Brameyer, S., et al. (2018), “A Single-Cell View of the BtsSR/YpdAB Pyruvate Sensing Network in Escherichia coli and Its Biological Relevance”, J Bacteriol, Vol. 200 No. 1, doi: 10.1128/JB.00536-17.

40. Weaver, J.W., Proshkin, S., Duan, W., Epshtein, V., Gowder, M., Bharati, B.K., Afanaseva, E., et al. (2023), “Control of transcription elongation and DNA repair by alarmone ppGpp”, Nat Struct Mol Biol, Vol. 30 No. 5, pp. 600–607, doi: 10.1038/s41594-023-00948-2.

41. World Health Organization. (2025), Global Antibiotic Resistance Surveillance Report 2025: WHO Global Antimicrobial Resistance and Use Surveillance System (GLASS), World Health Organization.

42. Yang, X., Liu, Y., Gan, J., Xiao, Z.X. and Cao, Y. (2022), “FitDock: protein-ligand docking by template fitting”, Brief Bioinform, Vol. 23 No. 3, doi: 10.1093/bib/bbac087.

43. Ye, J.B., Zeng, K., Li, X.B., Yang, J., Zhang, S. and Chen, Q. (2026), “Analysis of the efficacy of fosfomycin trometamol in preventing biofilm bacterial infection in double-J stents among diabetic patients and the factors associated with infection”, Int Urol Nephrol, doi: 10.1007/s11255-026-05080-w.

44. Zeng, T., Wang, Y., Zhu, Q., Xi, H., Liu, M. f, Liu, P., Bai, Y., et al. (2025), “Sub-inhibitory concentrations of fosfomycin enhance Staphylococcus aureus biofilm formation by a sarA-dependent mechanism”, Microbiol Spectr, Vol. 13 No. 9, p. e0152125, doi: 10.1128/spectrum.01521-25.

45. Zhang, H., Mu, R., Wang, Z., Peng, S., Yang, X.Y. and Qin, X. (2024), “Trans-Cinnamaldehyde Inhibition of Pyruvate Dehydrogenase: Effects on Streptococcus mutans Carbohydrate Metabolism”, J Proteome Res, Vol. 23 No. 8, pp. 3682–3695, doi: 10.1021/acs.jproteome.4c00382.

46. Zheng, D., Bergen, P.J., Landersdorfer, C.B. and Hirsch, E.B. (2022), “Differences in Fosfomycin Resistance Mechanisms between Pseudomonas aeruginosa and Enterobacterales”, Antimicrob Agents Chemother, Vol. 66 No. 2, p. e0144621, doi: 10.1128/AAC.01446-21.

