## Supplementary file for "Spicing up urinary tract infections: synergistic action of fosfomycin with trans-cinnamaldehyde - insight to the mode of action"

**Table S1.** Bacterial strains used in this study.

| No. | STRAIN | RELEVANT CHARACTERISTICS | REFERENCE |
| --- | --- | --- | --- |
|  | *Escherichia coli* |  |  |
| 1. | MG1655 | K-12 laboratory strain | (Blattner et al., 1997) |
| 2. | UTI89 | Human cystitis isolate; uropathogenic *E. coli* (UPEC) | (Fenlon et al., 2020) |
| 3. | CFT073 | Human pyelonephritis isolate; UPEC | (Welch et al., 2002) |
| 4. | KB077 | Clinical UPEC isolate | (Karczewska et al., 2023) |
| 5. | KB075 | Clinical UPEC isolate | (Karczewska et al., 2023) |
| 6. | KB076 | Clinical UPEC isolate | (Karczewska et al., 2023) |
| 7. | KB082 | Clinical UPEC isolate | (Karczewska et al., 2023) |
| 8. | KB083 | Clinical UPEC isolate | (Karczewska et al., 2023) |
| 9. | KB097 | Clinical UPEC isolate | (Karczewska et al., 2023) |
| 10. | KB047 | Clinical UPEC isolate | (Karczewska et al., 2023) |
| 11. | R4880 | Clinical FOS-resistant isolate carrying the *fos*A4 gene | (Mueller et al., 2019) |
| 12. | R82 | Clinical FOS-resistant isolate carrying the *fos*A3 gene | (Mueller et al., 2019) |
| 13. | N279 | Clinical FOS-resistant isolate; *fos*A1-*fos*A6 not detected | (Nordmann et al., 2019) |
| 14. | MG1655∆*aceE* | Deletion mutant lacking the E1 component (∆*aceE*::kanR*)* of the pyruvate dehydrogenase complex | Keio collection (Baba et al., 2006) |
| 15. | MG1655∆*aceF* | Deletion mutant lacking the E2 component (∆*aceF*::kanR) of the pyruvate dehydrogenase complex | Keio collection (Baba et al., 2006) |
| 16. | MG1655∆*ldcC* | Deletion mutant lacking constitutive lysine decarboxylase (∆*ldcC*::kanR*)* | Keio collection (Baba et al., 2006) |
| 17. | MG1655∆*poxB* | Deletion mutant lacking pyruvate oxidase (*poxB*::kanR) | Keio collection (Baba et al., 2006) |
| 18. | MG1655∆*lpdA* | Deletion mutant lacking pyruvate oxidase (*lpdA*::kanR) | Keio collection (Baba et al., 2006) |
| 19. | MG1655∆*btsSR* | Deletion mutant lacking the BtsSR two-component pyruvate-sensing system | (Vilhena et al., 2018) |
| 20. | MG1655∆*btsT* | Deletion mutant lacking BtsT, a high-affinity pyruvate transporter | (Kristoficova et al., 2018) |
| 21. | UTI89∆*aceE* | UTI89 derivative lacking the E1 component of the pyruvate dehydrogenase complex (∆*aceE*::FRT) | This study* |
| 22. | UTI89∆*aceF* | UTI89 derivative lacking the E2 component of the pyruvate dehydrogenase complex (∆*aceF*::FRT) | This study* |
| 22. | UTI89∆*lpdA* | UTI89 derivative lacking lipoamide dehydrogenase, the E3 component of the pyruvate dehydrogenase complex (∆*lpdA*::FRT) | This study* |

**UTI89 deletion mutants generated in this study were constructed using the λ Red recombination system. The kanamycin-resistance cassette was subsequently excised, leaving an FRT scar at the corresponding deletion locus, as indicated by Δ*gene*::FRT

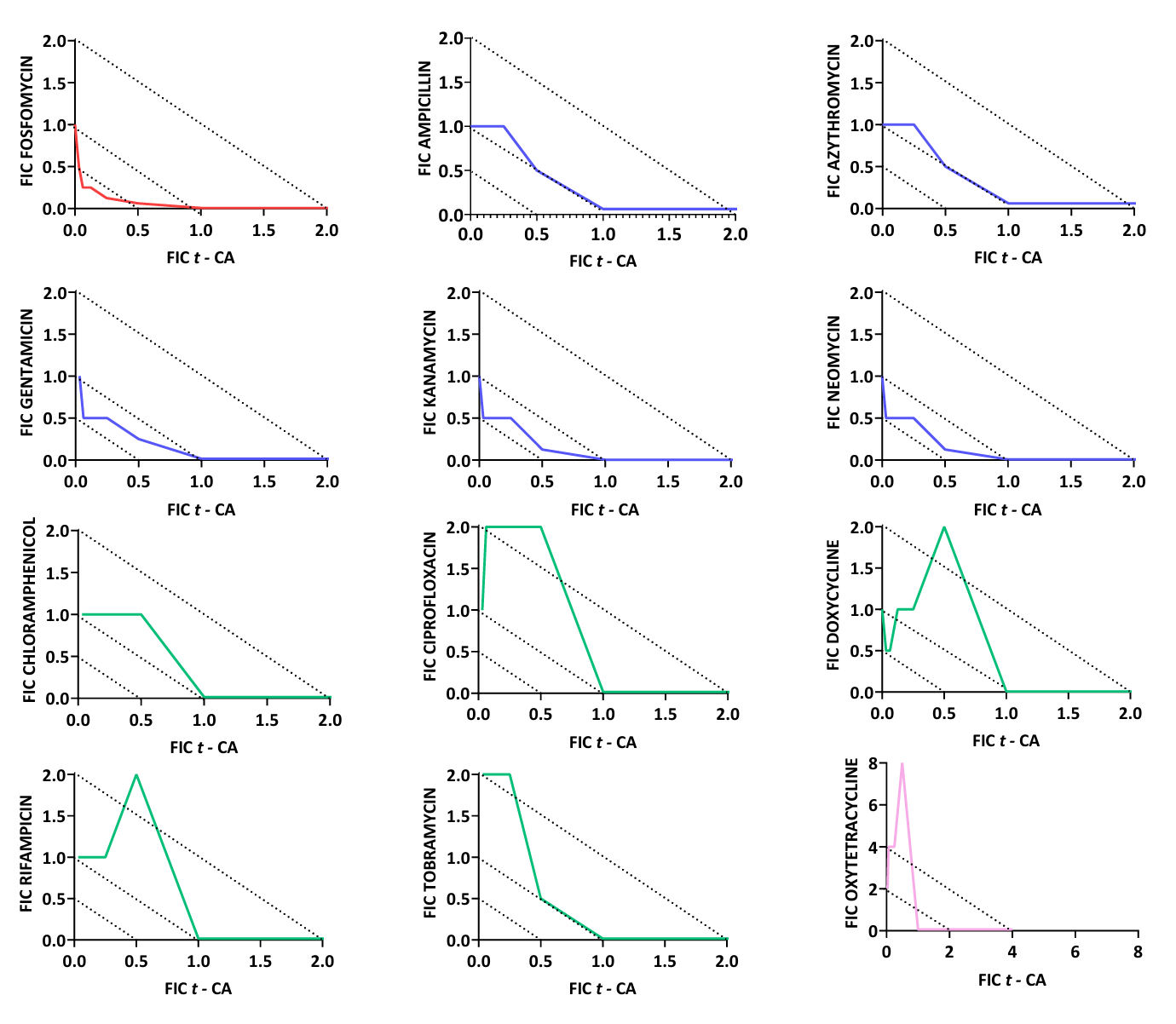

**Figure S1. Screening of interactions between t-CA and selected antibiotics.**
The *E. coli* K-12 strain MG1655 was used as the model strain. Isobologram/FIC plots showing the interaction between t-CA and representative antibiotics from different classes, including FOS, ampicillin, azithromycin, gentamicin, kanamycin, neomycin, chloramphenicol, ciprofloxacin, doxycycline, rifampicin, tobramycin, and oxytetracycline. The x-axis represents the fractional inhibitory concentration of t-CA, whereas the y-axis represents the fractional inhibitory concentration of the corresponding antibiotic. Assays were performed in at least three independent experiments. Dotted diagonal lines indicate interaction thresholds used for FICI interpretation: FICI < 0.5, synergism (red); 0.5–1.0, additive interaction (blue); 1.0–4.0, indifferent interaction (green); and >4.0, antagonism or unfavorable interaction (pink). Among the tested antibiotics, FOS showed the strongest synergistic interaction with t-CA, while most other combinations displayed additive or indifferent profiles. Oxytetracycline showed an unfavorable interaction profile under the tested conditions.

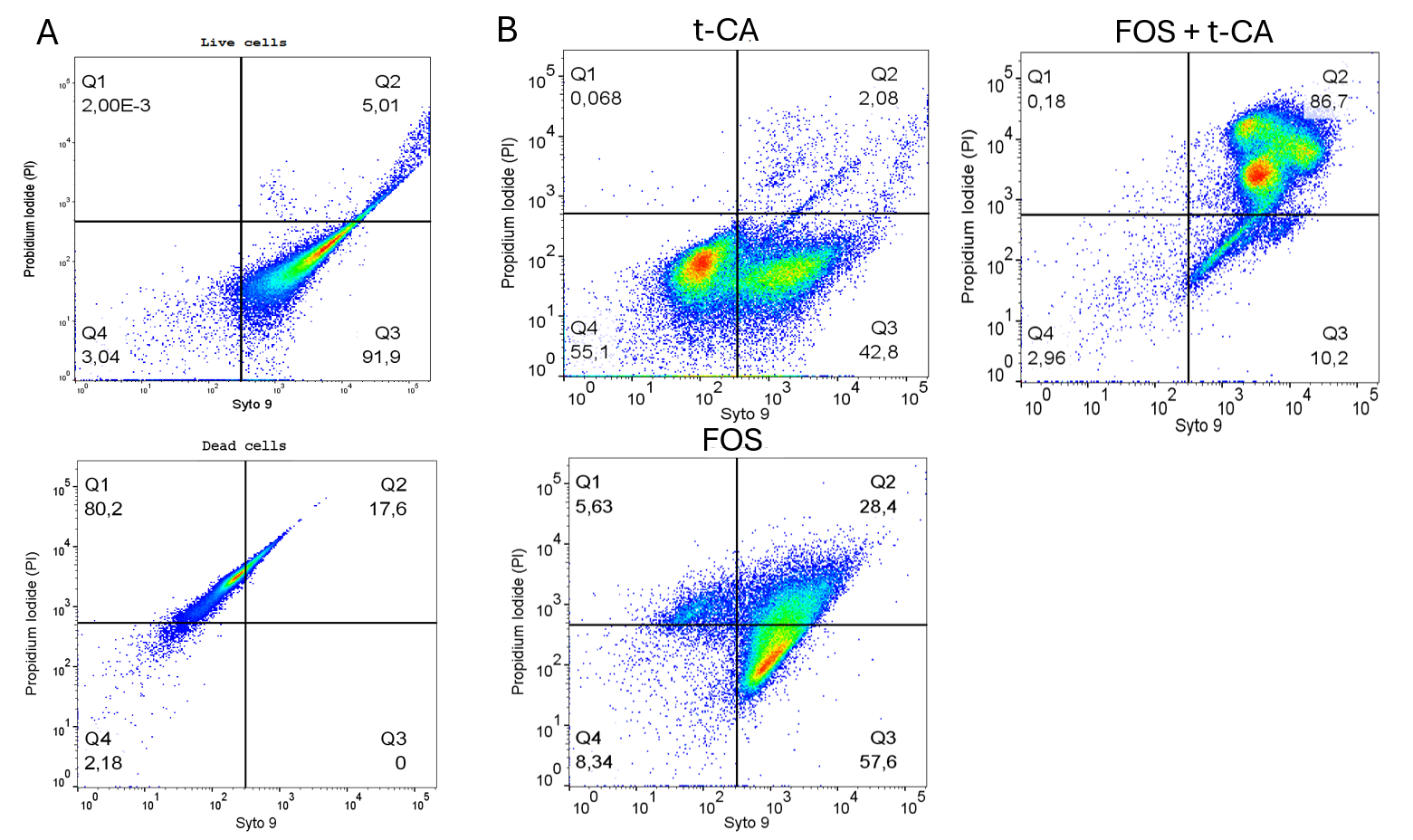

**Figure S2. Flow cytometric analysis of membrane integrity in UTI89 cells exposed to FOS, t-CA, or their combination.**
Representative SYTO 9/PI dot plots showing membrane integrity of UTI89 cells after 120 min treatment. Cultures were exposed to t-CA at 0.5×MIC, FOS at 1×MIC, or the t-CA/FOS combination at 0.5×MIC + 0.5×MIC. Untreated live cells and ethanol-killed cells were included as staining and gating controls. For each sample, 50,000 events were acquired. SYTO 9 fluorescence was used to identify bacterial cells, whereas PI fluorescence indicated loss of membrane integrity. Cells located in the SYTO 9-positive/PI-negative region were classified as viable, while PI-positive events were considered membrane-compromised or dead. Quadrant gates were set based on the live and ethanol-killed controls and applied uniformly to treated samples. The percentage of events in each quadrant is indicated on the plots.

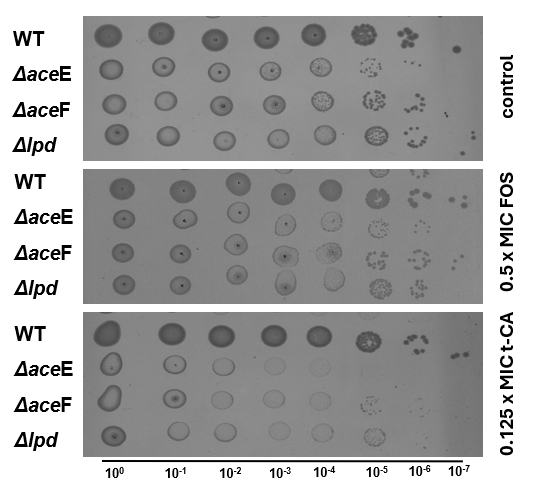

**Figure S3. Effect of the FOS or t-CA on *E. coli* UTI89 mutants impaired in pyruvate dehydrogenase activity.**
Serial dilution spot assay showing the growth of UTI89 WT and deletion mutants lacking individual components of the pyruvate dehydrogenase complex: Δ*aceE*, Δ*aceF*, and Δ*lpd*. Fresh grown bacterial cultures were serially diluted from 10⁰ to 10⁻⁷ and spotted onto LA control plates or plates containing FOS at 0.5×MIC or the t-CA at 0.125×MIC. Under control conditions and after exposure to FOS alone, all strains retained detectable growth across serial dilutions. In contrast, the t-CA/FOS combination markedly impaired the growth of PDH-deficient mutants, indicating that disruption of PDH-linked pyruvate metabolism sensitizes UPEC to the t-CA treatment.

References

Baba, T., Ara, T., Hasegawa, M., Takai, Y., Okumura, Y., Baba, M., Datsenko, K.A., Tomita, M., Wanner, B.L., Mori, H., 2006. Construction of Escherichia coli K-12 in-frame, single-gene knockout mutants: the Keio collection. Mol Syst Biol 2, 2006 0008. https://doi.org/10.1038/msb4100050

Blattner, F.R., Plunkett 3rd, G., Bloch, C.A., Perna, N.T., Burland, V., Riley, M., Collado-Vides, J., Glasner, J.D., Rode, C.K., Mayhew, G.F., Gregor, J., Davis, N.W., Kirkpatrick, H.A., Goeden, M.A., Rose, D.J., Mau, B., Shao, Y., 1997. The complete genome sequence of Escherichia coli K-12. Science (1979). 277, 1453–1462. https://doi.org/10.1126/science.277.5331.1453

Fenlon, S.N., Chee, Y.C., Chee, J.L.Y., Choy, Y.H., Khng, A.J., Liow, L.T., Mehershahi, K.S., Ruan, X., Turner, S.W., Yao, F., Chen, S.L., 2020. Sequencing of E. coli strain UTI89 on multiple sequencing platforms. BMC Res Notes 13, 487. https://doi.org/10.1186/s13104-020-05335-4

Karczewska, M., Strzelecki, P., Bogucka, K., Potrykus, K., Szalewska-Palasz, A., Nowicki, D., 2023. Increased Levels of (p)ppGpp Correlate with Virulence and Biofilm Formation, but Not with Growth, in Strains of Uropathogenic Escherichia coli. Int J Mol Sci 24. https://doi.org/10.3390/ijms24043315

Kristoficova, I., Vilhena, C., Behr, S., Jung, K., 2018. BtsT, a Novel and Specific Pyruvate/H(+) Symporter in Escherichia coli. J Bacteriol 200. https://doi.org/10.1128/JB.00599-17

Mueller, L., Cimen, C., Poirel, L., Descombes, M.C., Nordmann, P., 2019. Prevalence of fosfomycin resistance among ESBL-producing Escherichia coli isolates in the community, Switzerland. Eur J Clin Microbiol Infect Dis 38, 945–949. https://doi.org/10.1007/s10096-019-03531-0

Nordmann, P., Poirel, L., Mueller, L., 2019. Rapid Detection of Fosfomycin Resistance in Escherichia coli. J Clin Microbiol 57. https://doi.org/10.1128/JCM.01531-18

Vilhena, C., Kaganovitch, E., Shin, J.Y., Grunberger, A., Behr, S., Kristoficova, I., Brameyer, S., Kohlheyer, D., Jung, K., 2018. A Single-Cell View of the BtsSR/YpdAB Pyruvate Sensing Network in Escherichia coli and Its Biological Relevance. J Bacteriol 200. https://doi.org/10.1128/JB.00536-17

Welch, R.A., Burland, V., Plunkett 3rd, G., Redford, P., Roesch, P., Rasko, D., Buckles, E.L., Liou, S.R., Boutin, A., Hackett, J., Stroud, D., Mayhew, G.F., Rose, D.J., Zhou, S., Schwartz, D.C., Perna, N.T., Mobley, H.L., Donnenberg, M.S., Blattner, F.R., 2002. Extensive mosaic structure revealed by the complete genome sequence of uropathogenic Escherichia coli. Proc Natl Acad Sci U S A 99, 17020–17024. https://doi.org/10.1073/pnas.252529799
